# Plasma proteomics identifies an IL-6–associated SAA axis linked to muscle wasting and survival in patients with cancer cachexia

**DOI:** 10.64898/2026.06.11.730788

**Authors:** Jonas Sørensen, Christian T. Voldstedlund, Zakarias K. J. Ogueboule, Johanne Modvig, Cathrine Knøs, Joyceline Cuenco, Anna Leichter, Tabitha Cree, Anna H. Madsen, Christian S. Carl, Cecilie B. Lindqvist, Britanny Counts, Edmund Battey, Andrea Irazoki, Steffen H. Raun, Mona S. Ali, Emma Frank, Anne Bigot, Jan Christensen, Charlotte Suetta, Nicolai J. Wewer Albrechtsen, Kei Sakamoto, Teresa Zimmers, Richard Mills, Erik A. Richter, Geana P. Kurita, Seppo W. Langer, Julia S. Johansen, Inna M. Chen, Lykke Sylow

**Affiliations:** Department of Biomedical Sciences, Faculty of Health and Medical Sciences, University of Copenhagen, Copenhagen, Denmark; Department of Nephrology, Section of Palliative Medicine, Copenhagen University Hospital, Herlev-Gentofte, Denmark; Department of Nutrition, Exercise and Sports, Faculty of Science, University of Copenhagen, Copenhagen, Denmark; Novo Nordisk Foundation Center for Basic Metabolic Research, University of Copenhagen, Copenhagen, Denmark; Murdoch Children’s Research Institute, Royal Children’s Hospital, Melbourne, Victoria, Australia; Novo Nordisk Foundation Centre for Stem Cell Medicine (reNEW), Murdoch Children’s Research Institute, Melbourne, Victoria, Australia; Department of Paediatrics, School of Biomedical Sciences, University of Melbourne, Melbourne, Victoria, Australia; Department of Cell, Developmental and Cancer Biology, Oregn Health & Science University, Portland, OR, USA; Sorbonne Université, Inserm, Institut de Myologie, Centre de Recherche en Myologie, Paris, France; Department of Public Health, Faculty of Health and Medical Sciences University of Copenhagen, Copenhagen, Denmark; Department of Occupational Therapy and Physiotherapy, Centre of Head and Orthopaedic Surgery, Copenhagen University Hospital – Rigshospitalet, Copenhagen, Denmark; Department of Clinical Medicine, Faculty of Health and Medical Sciences, University of Copenhagen, Copenhagen, Denmark; Department of Geriatric and Palliative Medicine, Bispebjerg and Frederiksberg Hospital, Copenhagen, Denmark; Department of Clinical Biochemistry Copenhagen University Hospital, Bispebjerg, Copenhagen, Denmark; Portland Veterans Administration Medical Center, Portland, OR, USA; Knight Cancer Institute, Oregn Health & Science University, Portland, OR, USA; Brendon-Colson Center for Pancreatic Care, Oregon Health & Science University, Portland, OR, USA; Department of Anaesthesiology, Multidisciplinary Pain Centre, Pain and Respiratory Support, Copenhagen University Hospital – Rigshospitalet, Copenhagen, Denmark; St. Luke’s Academy of Palliative Medicine, St. Luke’s Foundation, Hellerup, Denmark; Department of Oncology, Copenhagen University Hospital - Rigshospitalet, Copenhagen, Denmark; Department of Oncology, Copenhagen University Hospital - Herlev and Gentofte, Herlev, Denmark; Department of Medicine, Copenhagen University Hospital - Herlev and Gentofte, Herlev, Denmark

**Author notes:** These authors contributed equally to this work (Jonas Sørensen, Christian T. Voldstedlund, Zakarias K. J. Ogueboule).

**Keywords:** cancer cachexia, non-small cell lung cancer, pancreatic cancer, muscle wasting, fat wasting, plasma, proteomics, body composition, serum amyloid A1 (SAA1) and A2 (SAA2), IL-6, human myotubes

## Abstract

Cachexia affects nearly half of patients with lung- and pancreatic cancer and worsens prognosis, yet factors driving muscle wasting remain poorly defined. We performed longitudinal clinical and plasma proteomic profiling of 67 patients with non-small cell lung cancer, with and without cachexia, during first-line treatment. At diagnosis, cachectic patients showed 128 upregulated and 67 downregulated plasma proteins relative to non-cachectic patients. Longitudinal body composition, physical performance, clinical outcome, and nutritional risk assessments revealed distinct fat- and muscle-wasting trajectories: 71 proteins tracked with fat loss, 92 with muscle loss, and 177 with concurrent muscle and weight loss. Patients with concurrent loss had 68% shorter median overall survival than those who preserved mass (239 vs. 740 days).

Among proteins associated with both muscle loss and adverse outcomes, we prioritized eight candidates and screened their recombinant forms, where serum amyloid A (SAA) produced a 47% reduction in myotube width and impaired contractile function in human 2D and 3D myotubes. In an independent cohort of 146 patients with advanced pancreatic cancer, pharmacological blockade of IL-6 signaling with tocilizumab normalized circulating SAA levels, establishing a cancer-driven IL-6–SAA axis that promotes muscle wasting. In both lung and pancreatic cancer, elevated SAA was associated with reduced overall survival. Together, these findings identify SAA1 and SAA2 as IL-6-driven, cachexia-associated factors that impair muscle mass and function and predict survival, nominating the IL-6–SAA axis as a therapeutic target for cancer cachexia.

**Graphical abstract:** 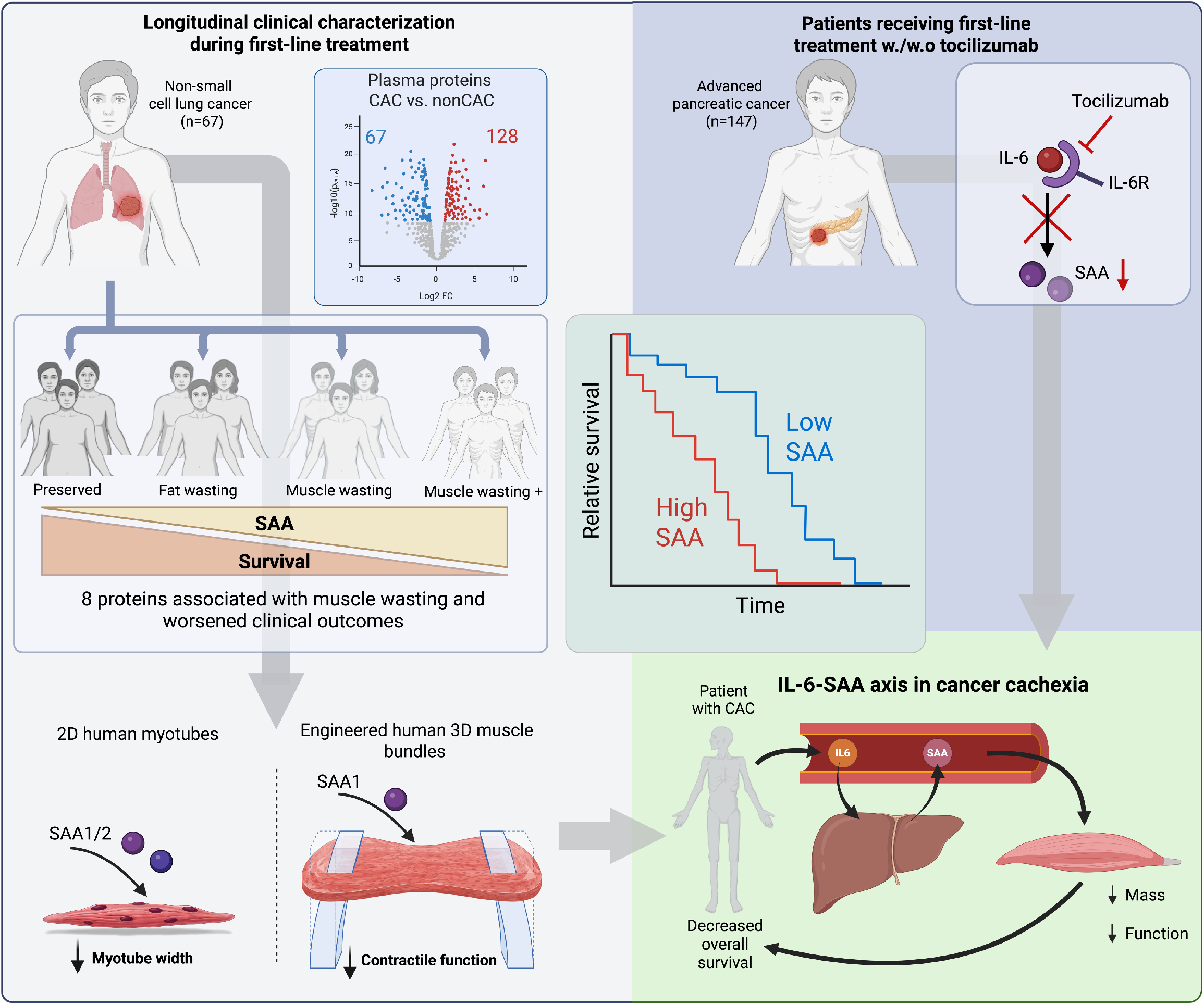

## INTRODUCTION

Cancer-associated cachexia (CAC) is a severe wasting syndrome and a major contributor to mortality^1,2^, yet the underlying pathophysiology remains poorly understood and is particularly understudied in humans. Among the 2.3 million individuals diagnosed with lung cancer and 500,000 with pancreatic cancer (PC) globally each year, more than half develop CAC^3,4^, characterized by anorexia and a rapid loss of muscle and fat mass^5,6^. Lacking therapies approved by the European Medicines Agency (EMA) or the U.S. Food and Drug Administration (FDA), cachexia impairs quality of life, increases treatment toxicity, and substantially compromises survival^7–10^, underscoring the urgent need to identify molecular determinants that drive or predict its onset and ultimately guide future interventions.

Advances in mass spectrometry (MS)-based proteomics have transformed biomedical research by enabling unbiased, high-throughput protein quantification across large clinical cohorts^11^. This phenotyping facilitates the discovery of novel disease mechanisms and actionable biomarkers^12–15^. Human studies encompassing biomarkers with longitudinal changes in body composition, clinical trajectories, and patient-reported measures remain scarce; therefore, there is an urgent need to align systemic biomarkers with clinical changes. Such integration is essential to elucidate cachexia pathobiology at the plasma proteomic level. An example is the identification of circulating GDF15 as a candidate biomarker of CAC and the subsequent clinical efficacy of the GDF15-neutralizing antibody ponsegromab in a randomized phase II trial, demonstrating that circulating proteins can be therapeutically targeted in CAC^16–18^.

Our study had four primary aims: (1) to identify individuals with CAC at diagnosis and assess associated clinical burdens; (2) to map plasma proteomic alterations driving distinct muscle and fat wasting trajectories; (3) to functionally validate the direct role of candidate circulating proteins in human myotubes; and (4) to delineate the upstream regulatory dependency of these proteins using targeted pharmacological inhibition in an independent clinical cohort.

We conducted a comprehensive clinical assessment and MS-based plasma proteomic analysis to identify circulating proteins linked to cachexia in patients with advanced-stage NSCLC, both at the diagnosis and during the first 12 weeks of first-line treatment, with subsequent annotation and functional validation of candidate proteins in human myotubes. We identified 195 plasma proteins that were differentially regulated in NSCLC patients with CAC compared with patients without CAC. By integrating longitudinal functional assessments, body imaging, patient-reported nutritional risk, and blood sampling with recent technological advances in deep plasma proteomic phenotyping, we identified 95 plasma proteins associated with fat wasting, 92 with muscle wasting, and 177 associated with concurrent muscle wasting and weight loss during treatment. Pharmacological suppression of serum amyloid A (SAA) levels following IL-6 inhibition in patients with advanced PC, together with our clinically anchored unbiased selection workflow, identified an IL-6-SAA axis that potentially mediates muscle wasting in patients with cancer cachexia. Altogether, these findings provide new insights into the CAC pathobiology and highlight novel potential therapeutic targets.

## RESULTS

### Characteristics of study participants

To identify patients with CAC at diagnosis, 107 patients with advanced-stage NSCLC were assessed for eligibility, and 75 were included at diagnosis in this longitudinal observational study, following the inclusion and assessment flow illustrated in Figure 1. Eight patients were lost to follow-up before week 6, leaving 67 patients for the longitudinal analyses. The cohort was evenly balanced by sex, with a mean age of 65 years. Of the 67 included patients, 36 had CAC and 16 had pre-cachexia (preCAC) (Table 1); 56% of CAC patients were male. Of the 67 patients, 3 received platinum-based chemotherapy or vinorelbine, 22 received an immune checkpoint inhibitor, 32 received a combination of these, and 10 were treated with a tyrosine-kinase inhibitor (Table 1). Apart from Eastern Cooperative Oncology Group Performance Status (PS) (p=0.016), no sociodemographic or disease-specific differences were observed between the 75 included and 67 analyzed patients (Table 1). The risk of attrition bias from loss to follow-up was therefore low, and our NSCLC cohort aligns with characteristics reported in other studies^19^.

**Figure 1.**
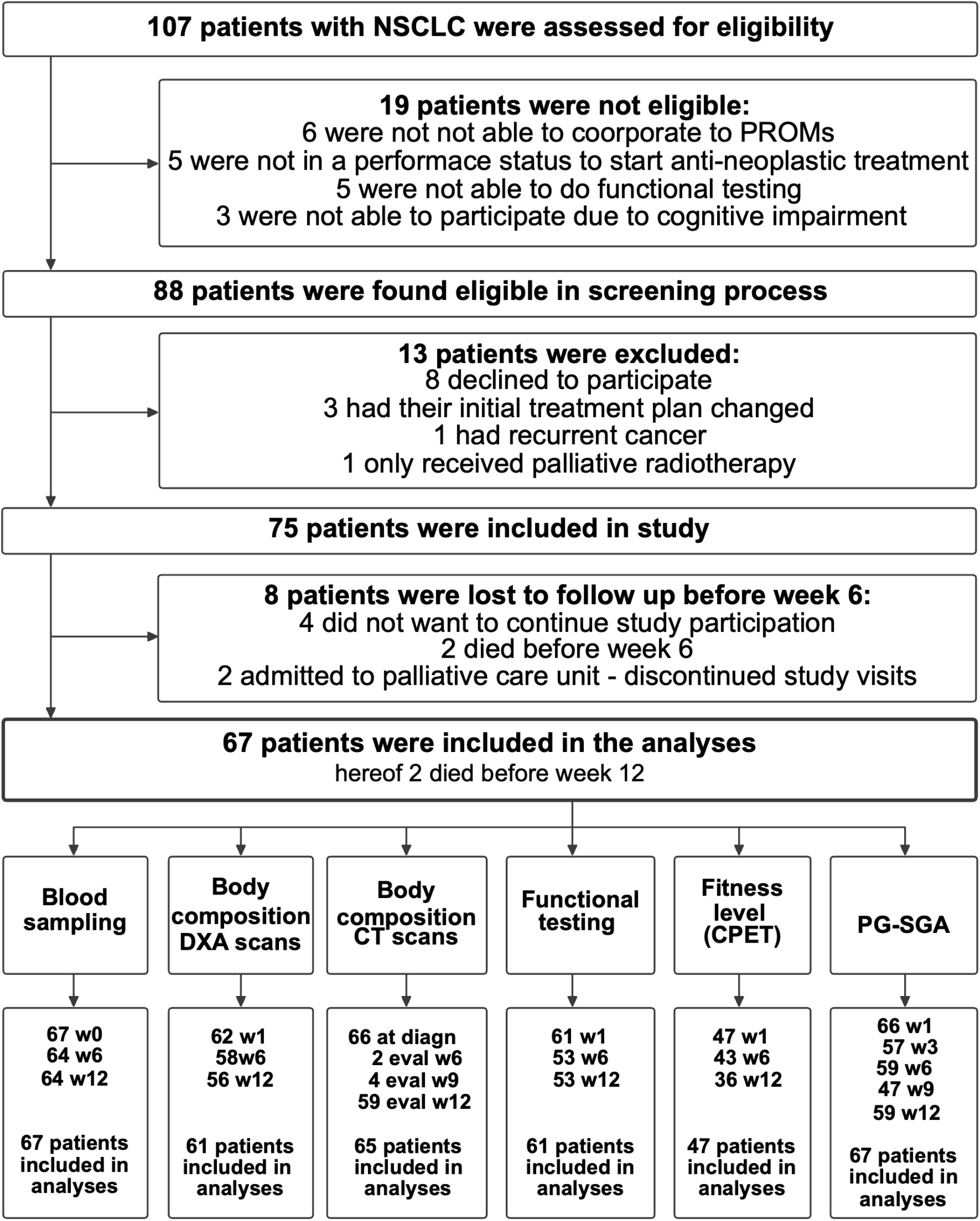
Study profile: inclusion and assessment flow. Inclusion and assessment flow in the study. Of 107 patients with non-small cell lung cancer (NSCLC) assessed for eligibility, 75 were included. Longitudinal data were obtained for blood sampling (67 patients), DXA-derived body composition (61), CT-derived body composition (65), functional testing (61), fitness level by CPET (47), and PG-SGA patient-reported outcomes (67); see the bottom row of the figure for numbers at each timepoint. DXA=dual-energy X-ray absorptiometry. CT=computed tomography. CPET=cardiopulmonary exercise test. PG-SGA=patient-generated subjective global assessment. w1=week 1 after NSCLC diagnosis; w3=week 3; w6=week 6; w9=week 9; w12=week 12.

**Table 1.** NSCLC patient characteristics at diagnosis; hospitalization and treatment-delaying toxicity during treatment. Data are n (%) or mean (SD), unless otherwise noted; percentages use the number of patients in each row as the denominator. The right-hand column reports socio-demographic and disease-specific data for all 75 patients included in the study; the middle column reports the subgroup of 67 patients included in the longitudinal analyses. *Four patients reported their weight one month before diagnosis; this was used for weight-loss/cachexia stratification at diagnosis. **Simple logistic regression with Holm’s correction found only PS significantly different between the groups (p=0.016). PS=Eastern Cooperative Oncology Group Performance Score. PD-L1=programmed death-ligand 1. FEV1=forced expiratory volume in 1 second. CAC=cachexia. preCAC=pre-cachexia. nonCAC=no cachexia. BMI=body mass index. CRP=C-reactive protein. IL-6=interleukin-6. CT=computed tomography. SMI=skeletal muscle index. L3/CT=lumbar vertebral level 3 on CT. IQR=interquartile range.

| Variable | Patients |  |
| --- | --- | --- |
|  | Included in the analyses | Included in the study |
| **Sex** | N=67 | N=75 |
| Male, n (%) | 33 (49) | 36 (48) |
| Female, n (%) | 34 (51) | 39 (52) |
| **Age** | N=67 | N=75 |
| Years, mean(SD) | 65 (10.3) | 65 (10.1) |
| Min-max | 39-81 | 39-83 |
| **Cachexia status, at time of diagnosis\*** | N=67 | n=70 |
| CAC, n (%) | 36 (54) | 39 (56) |
| >5% weight loss last 6 months | 27 (40) | 30 (43) |
| >2% weight loss last 6 months + BMI <20 | 3 (5) | 3 (4) |
| >2% weight loss last 6 months + sarcopenia, SMI defined | 6 (9) | 6 (9) |
| preCAC, n (%) | 16 (24) | 16 (23) |
| 1% < weight loss last 6 months ≤ 5 %, BMI ≥ 20, no sarcopenia |  |  |
| nonCAC, n (%) | 15 (22) | 15 (21) |
| **Primary treatment plan** | N=67 | N=75 |
| Platinum-based chemotherapy or vinorelbine, n (%) | 3 (4) | 5 (7) |
| Immune check-point inhibitor, n (%) | 22 (33) | 26 (35) |
| Combined chemotherapy / immune check-point inhibitor, n (%) | 32 (48) | 34 (45) |
| Tyrosine kinase inhibitor, n (%) | 10 (15) | 10 (13) |
| **Eastern Cooperative Oncology Group Performance Score (PS)\*\*** | N=67 | N=75 |
| 0, n (%) | 28 (42) | 28 (37) |
| 1, n (%) | 37 (55) | 42 (56) |
| 2, n (%) | 2 (3) | 5 (7) |
| **Histology** | N=67 | N=75 |
| Adenocarcinoma, n (%) | 53 (79) | 60 (80) |
| Squamous cell carcinoma, n (%) | 10 (15) | 10 (13) |
| Other, n (%) | 4 (6) | 5 (7) |
| **PD-L1 status** | N=67 | N=75 |
| Positive >50%, n (%) | 28 (42) | 32 (43) |
| Positive >1 %, n (%) | 22 (33) | 23 (31) |
| Negative, n (%) | 17 (25) | 20 (27) |
| **Age-adjusted Charlson Comorbidity Index (aCCI)** | N=67 | N=75 |
| Mean (SD) | 9.7 (2.0) | 9.8 (1.9) |
| Min-max | 6-15 | 6-15 |
| **Current pharmacological treatment** | N=67 | N=75 |
| Daily oral/sc antidiabetics, n (%) | 4 (6) | 5 (7) |
| Daily use of corticosteroids, n (%) | 4 (6) | 4 (5) |
| Daily use of beta-blockers, n (%) | 9 (13) | 9 (12) |
| **Smoking & pulmonary status** |  |  |
| Current smokers, n (%) of reg. | 14 (22) 65 | 15 (22) 66 |
| Smoking pack year, mean (SD), min-max, of reg. | 33.5 (22.0),<br>0-100, 66 | 33.8 (22),<br>0-100, 67 |
| FEV1 % of expected, mean (SD), min-max, of reg. | 68.8 (19.9),<br>27-121, 66 | 68.7 (20),<br>27-121, 67 |
| **Sarcopenia status** | N=65 |  |
| According to SMI from L3/CT: |  |  |
| Sarcopenia, n (%) | 41 (63) |  |
| Non sarcopenia, n (%) | 24 (37) |  |
| **BMI** | N=67 |  |
| Mean (SD) | 24.4 (4.5) |  |
| Min-max | 16.4-41.4 |  |
| BMI < 20 kg/m<sup>2</sup>, n (%) | 11 (16) |  |
| **CRP (mg/L)** | N=61 |  |
| at diagnosis, median | 25 |  |
| Min-max | 1-207 |  |
| < 10, n (%) | 17 (28) |  |
| 10 - 49, n (%) | 22 (36) |  |
| ≥ 50, n (%) | 22 (36) |  |
| **Albumin (g/L)** | N = 67 |  |
| at diagnosis, mean (min-max) | 34 (20-42) |  |
| **IL-6 (pg/mL)**<br>at diagnosis, median (min-max)<br>CAC (n=36)<br>preCAC (n=16)<br>nonCAC (n=15) | N=67<br>10.6 (1.5-237.1)<br>18.9 (1.5-237.1)<br>10.1 (1.5-90.8)<br>4.75 (1.5-18.1) |  |
| **Hospitalization, related to cancer or treatment hereof**<br>Yes, n (%)<br>No, n (%) | N=67<br>26 (39)<br>41 (61) |  |
| **Treatment delaying toxicity**<br>Yes, n (%)<br>No, n (%) | N=67<br>27 (40)<br>40 (60) |  |
| **Data collection, selected assessments**<br>Diagnosis (days *before* inclusion)<br>(*median; IQR (min-max)*)<br>Baseline CT scan (days *before* inclusion)<br>(*median; IQR (min-max)*)<br>Functional testing, week 1 (days *after* inclusion)<br>(*median; IQR (min-max)*)<br>Evaluating CT scan (days *after* inclusion)<br>(*median; IQR (min-max)*) | N=67<br>8; 4 (1-49)<br>N=67<br>24; 9 (14-44)<br>N = 62<br>4; 5 (0-14)<br>N=66<br>75; 8.5 (47-124) |  |

### CAC at NSCLC diagnosis associates with nutritional risk, hospitalization, and treatment-delaying toxicity across 12 weeks of first-line treatment

Following baseline CAC identification, we evaluated its association with nutritional risk, hospitalization, and treatment-limiting toxicities to define the clinical burden of CAC throughout treatment. Low appetite and decreased functional capacity are clinical signs of CAC^20^, which were captured in the patient-generated subjective global assessment (PG-SGA) score during treatment^21^. Mean PG-SGA scores remained stable throughout first-line treatment (70–98% response rate), with most patients scoring below the clinical threshold of 6, indicating nutritional risk^22^ (Figure S1A). To investigate patient-reported nutritional risk in patients with CAC at diagnosis, we employed linear mixed modeling and found that baseline CAC was significantly associated with longitudinal PG-SGA scores (p=0.0031, Figure S1B). These findings highlight CAC at diagnosis as a key factor of nutritional risk during treatment, despite individualized and tailored nutritional counseling (see Method section).

CAC imposes a profound individual, societal, and treatment burden, manifesting in increased hospitalization and heightened treatment-related toxicity^20,23^. Accordingly, we found that patients with nonCAC had a mean hospitalization time of 1.9 days during treatment, while patients with preCAC or CAC at diagnosis averaged 6.9 and 5.6 days, respectively (Figure S1C). Over 12 weeks, treatment-delaying toxicities occurred in 44.4% and 43.8% of patients with CAC and preCAC, respectively, compared to 26.7% in nonCAC patients, highlighting a markedly higher clinical burden (Figure S1D). This yields a need for early identification and proactive monitoring to optimize clinical outcomes in this high-risk population.

### Profiling body composition and weight loss during treatment reveals distinct wasting phenotype trajectories correlating with key clinical variables

Body composition undergoes dramatic alterations in response to cancer. Post-diagnosis weight loss is a critical negative prognostic factor in NSCLC, driven by distinct wasting of skeletal muscle, adipose tissue, or both^8^. While such phenotypes are recognized in pancreatic cancer^24^, the heterogeneity and individual wasting trajectories in NSCLC remain uncharacterized. To address this gap, we integrated computed tomography (CT)-derived skeletal muscle (SMI) and subcutaneous adipose tissue (SATI) indices with weight loss data to define individual wasting trajectories. We stratified the 67 patients by CAC status at diagnosis^25^ and by wasting phenotype trajectories. Patients who maintained or gained weight without muscle or fat loss were categorized as “Preserved,” while others were stratified into three distinct wasting phenotypes: Fat Wasting (FW), Muscle Wasting without weight loss (MW), and Muscle Wasting with concurrent weight loss (MW+). We did not observe associations (Figure 2A) between CAC at diagnosis and the 12 weeks progression of the distinct wasting phenotypes, suggesting that CAC at diagnosis cannot predict the wasting trajectories during first-line NSCLC treatment. Details on CT- and dual-energy X-ray absorptiometry (DXA)-derived body compositions, weight, and CT-derived wasting phenotypes for all patients during treatment can be found in Table S1.

**Figure 2.**
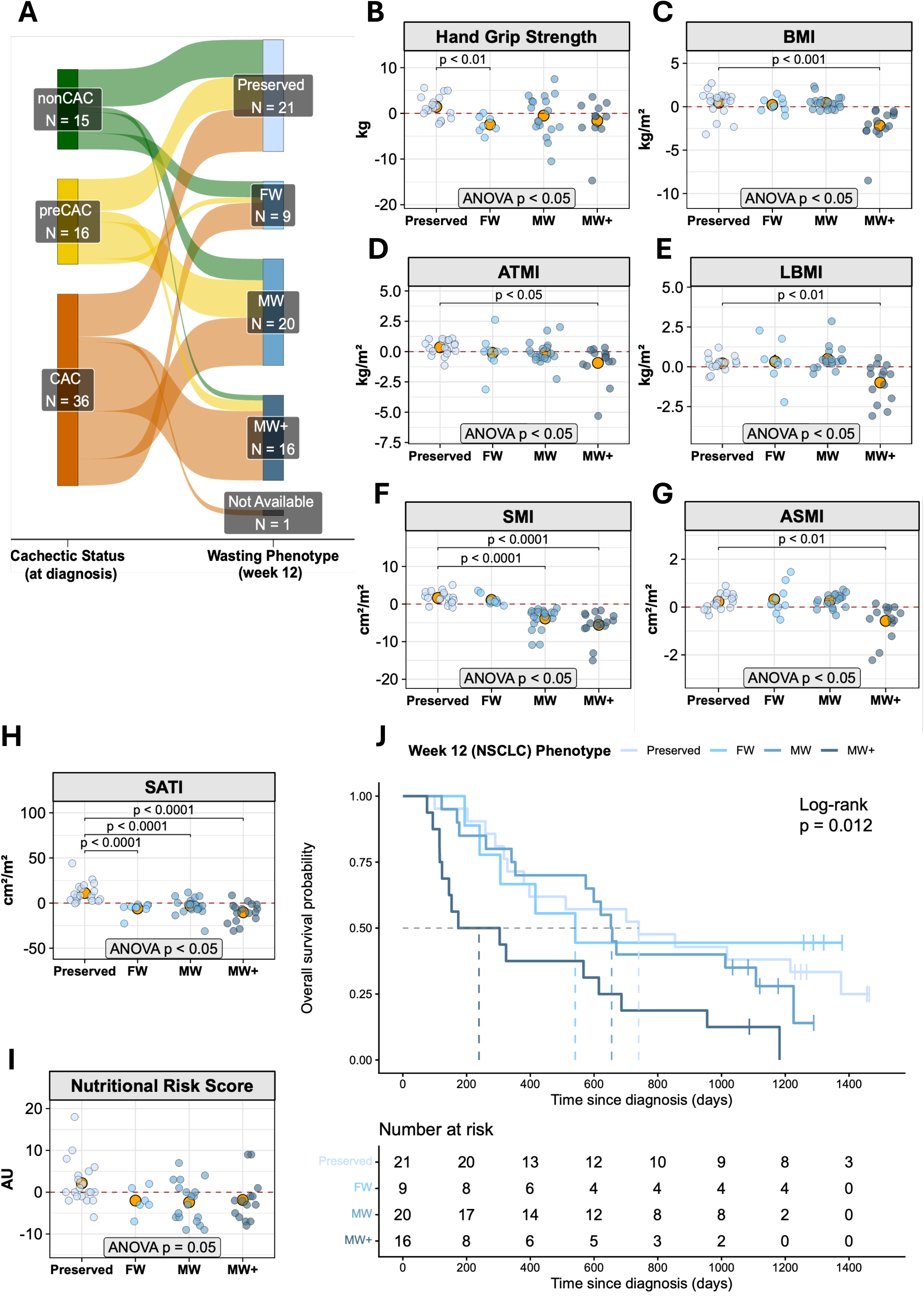
Profiling weight loss across treatment reveals specific wasting phenotypes and marked skeletal muscle wasting. **A:** Sankey diagram showing CAC status at diagnosis and its contribution to three wasting-phenotype trajectories over 12 weeks of first-line treatment. Band width is proportional to patient number, so the widest bands represent the dominant clinical phenotypes. Wasting phenotypes were defined by CT-derived muscle and weight loss: Preserved (no wasting; <2.5% loss of both SMI and SATI), fat wasting (FW; SATI loss >2.5%), muscle wasting without weight loss (MW), and muscle wasting with weight loss (MW+; total body weight loss >2.5% distinguishes MW from MW+). One patient was lost to follow-up CT at week 12; one patient with a technical CT error was classified as Preserved based on weight gain and unchanged DXA measures. nonCAC=no cachexia. preCAC=pre-cachexia. CAC=cachexia. **B-I:** Changes in clinical variables from diagnosis to week 12 across wasting phenotypes: hand grip strength (B), BMI (C), adipose tissue mass index (ATMI, D), lean body mass index (LBMI, E), skeletal muscle index (SMI, F), appendicular skeletal muscle index (ASMI, G), subcutaneous adipose tissue index (SATI, H), and PG-SGA nutritional risk score (I). Semi-transparent dots represent individual patients; orange dots show the mean. Panels shown had a significant one-way ANOVA across phenotypes (p<0.05); brackets show unpaired t-test pairwise comparisons with significance levels. kg=kilograms. BMI=body mass index. ATMI=adipose tissue mass index. LBMI=lean body mass index. SMI=skeletal muscle index. ASMI=appendicular skeletal muscle index. SATI=subcutaneous adipose tissue index. AU=arbitrary unit. PG-SGA=patient-generated subjective global assessment. Preserved=no wasting. FW=fat wasting. MW=muscle wasting without weight loss. MW+=muscle wasting with weight loss. **J:** Overall survival by week-12 wasting phenotype: Preserved (n=21), FW (n=9), MW (n=20), MW+ (n=16); one patient who died before the week-12 assessment could not be classified and was excluded (n=66 total). Survival was calculated from diagnosis to death from any cause; data were censored at the September 3, 2026 cutoff (vertical tick marks). Numbers at risk are shown below the plot. Groups were compared by log-rank test.

Beyond disease- and patient-related factors driving muscle wasting, a direct wasting effect of anti-cancer treatment cannot be ruled out ^26–29^. However, our secondary analysis of CAC status across the four treatment groups at diagnosis (Table 1) revealed no clear association between regimen and phenotypic wasting trajectories (Figure S1E).

Having defined distinct wasting phenotypes, we investigated their associations with key clinical variables during first-line NSCLC treatment. Phenotype-specific analysis revealed that handgrip strength was impaired in FW patients relative to the Preserved group (Figure 2B). BMI (Figure 2C), adipose tissue mass index (ATMI) (Figure 2D) and lean body mass index (LBMI) (Figure 2E), SMI (Figure 2F), appendicular skeletal muscle index (ASMI) (Figure 2G) and SATI (Figure 2H) were all reduced in patients with MW+ compared to Preserved. The reduction of SMI and SATI in MW patients relative to the Preserved cohort validates our CT-derived classification as a robust, clinically accessible tool. No differences were observed between the wasting phenotypes in fitness (Figure S2A), sit-to-stand performance (Figure S2 B), skeletal muscle radio attenuation (Figure S2C), the nutritional risk score (Figure 2I) or in the biochemical variables hemoglobin (Hgb) (Figure S2D), HbA1c (Figure S2E), albumin (Figure S2F) and c-reactive protein (CRP) (Figure S2G). Although underpowered to stratify survival by wasting trajectory, Kaplan-Meier analysis indicated a reduced survival in MW+ NSCLC patients who displayed 68% shorter median overall survival (239 days) compared patients in the Preserved group (740 days) (Figure 2J).

Collectively, integrating baseline CAC status with longitudinal body composition profiling reveals the inherent complexity and heterogeneity of wasting trajectories in advanced NSCLC. These phenotypes effectively differentiate changes in clinical variables and possibly survival, establishing their clinical relevance and providing a framework for uncovering the molecular signatures of these distinct trajectories.

### Plasma proteomic profiling reveals molecular signatures associated with CAC at diagnosis and identifies proteins linked to distinct wasting phenotype trajectories

Following CAC categorization and the identification of distinct wasting phenotypes, we investigated the plasma proteome to uncover molecular signatures associated with CAC and specific phenotype trajectories.

At diagnosis, plasma proteomic profiling of 3,078 proteins identified 195 proteins differentially abundant in CAC relative to nonCAC patients (128 upregulated; 67 downregulated; Figure 3A). Of these, 19 proteins (SPINK1, SMG1, IGHV4-34, PEAK3, CLN5, ARHGAP23, PLA2G2A, TNS1, IGHD, NANP, NCK1, CRP, PHF6, NUAK2, TGM2, IGHE, SAA2, KANK1 and AOAOJ9YY99) were upregulated (Log2FC > 1.0), whereas four proteins (LNPK, PLA2G1B, GCLC and MANSC1) were downregulated (Log2FC < −1.0). Principal Component Analysis (PCA) revealed no distinct separation between groups (Figure S2H), suggesting that global plasma proteome alterations do not distinguish CAC from nonCAC patients. However, enrichment analysis (Figure S2I) revealed that CAC patients exhibited a highly inflammatory signature, as indicated by significant enrichment of adaptive “immune response”, “response to bacterium” and several immunoglobin related immune pathways, in line with the classification of cachexia as a systemic inflammatory disease^18,30^.

**Figure 3.**
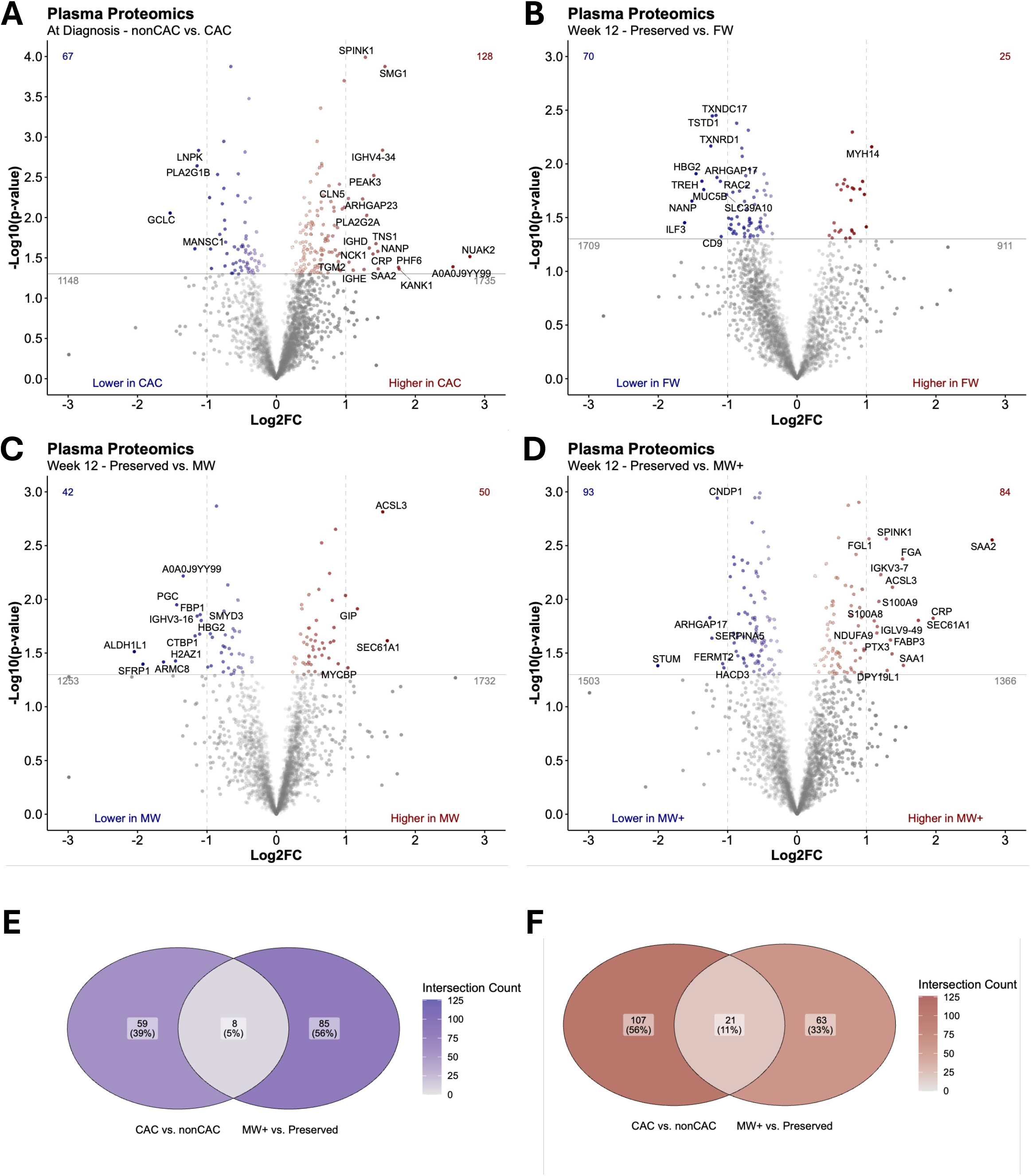
Plasma proteome at diagnosis and during 12 weeks of first-line NSCLC treatment. **A-D:** Volcano plots of differential plasma proteome abundance: nonCAC vs. CAC at diagnosis (A); Preserved vs. FW (B), vs. MW (C), and vs. MW+ (D) at week 12. **E-F:** Venn diagrams showing the overlap between the nonCAC-vs-CAC (diagnosis) and Preserved- vs-MW+ (week 12) comparisons, for downregulated (E) and upregulated (F) proteins. P values were calculated by unpaired Welch’s t-test with permutation-based recalibration. Preserved=no wasting. FW=fat wasting. MW=muscle wasting without weight loss. MW+=muscle wasting with weight loss. nonCAC=no cachexia. CAC=cachexia. Log2FC=log2 fold change.

Having determined plasma proteomic signatures associated with CAC at diagnosis, we next sought to identify plasma proteins linked to the three distinct wasting phenotype trajectories during treatment. We observed divergence in the plasma proteome when comparing Preserved vs. FW (25 up/70 down; Figure 3B) and MW (42 up/50 Down; Figure 3C), and MW+ (84 up/93 down; Figure 3D). Similar to the comparison between nonCAC and CAC, these phenotypic differences were not reflected in the PCA plot (Figure S2J,K,L), suggesting that increasing phenotypic muscle wasting severity is associated with more confined alterations in protein abundance rather than broad global proteomic remodeling. Intriguingly, when further analyzing the Preserved vs. MW+ (Figure 3D) we observed 17 proteins significantly upregulated (p<0.05) and with Log2FC>1.0 (FGL1, SPINK1, FGA, ACSL3, S100A9, IGKV3-7, S100A8, SEC61A1, IGLV9-49, CRP, FABP3, NDUFA9, HMBS, PTX3, SAA1, SAA2, and DPY19L1). These proteins all stand out with the potential of playing crucial roles in a highly cachectic patient phenotype with substantial muscle wasting and weight loss, known to be associated with reduced survival in patients with NSCLC^8^. Among these MW+-associated proteins, 5% were co-downregulated (Figure 3E) and 11% co-upregulated (Figure 3F) in CAC patients, indicating predominantly distinct mechanisms with relatively modest proteomic overlap (supporting/supplemental data). Over-representation analysis of the plasma proteome from MW+ patients, exhibiting muscle wasting and weight loss, revealed enrichment of inflammatory acute phase response (Figure S2M). Secondary analysis of the treatment groups (Figure S2N) indicated that the observed plasma proteomic variations were independent of anti-cancer therapy. Altogether, our MS-based plasma proteomic analyses revealed distinct molecular signatures associated with CAC at diagnosis and with specific wasting phenotype trajectories during the first 12 weeks of NSCLC treatment and subsequent survival, providing robust evidence that identify systemic inflammation as a primary therapeutic target to treat cachexia.

### Target selection workflow identifies eight plasma proteins as proteomic signatures of skeletal muscle wasting in NSCLC patients during treatment

Skeletal muscle wasting has been associated with poor prognosis, even in weight stable patients^9,31^. We therefore focused on plasma proteomic changes associated with muscle wasting trajectories, with and without concurrent weight loss, being MW and MW+. To identify proteins linked to patient outcomes, we developed a targeted selection workflow integrating plasma proteins significantly regulated in CAC at diagnosis, proteins associated with muscle wasting trajectories, and proteins correlated with key clinical variables (Figure 4A). Twelve proteins met at least two criteria and were identified as potential contributors to muscle wasting: CLN5, A0A0J9YY99, SMG1, CNDP1, FGA, FGL1, S100A8, S100A9, SAA1, CRP, SAA2, and SPINK1. A0A0J9YY99 was excluded because its function remains poorly characterized, with only automated, unverified annotation available. To define their collective dynamics, we modeled a correlation network at diagnosis and week 12. Eight proteins (S100A8, S100A9, CRP, FGL1, SPINK1, CLN5, SAA1 and SAA2) clustered and correlated with key clinical variables both at diagnosis (Figure 4B) and after treatment (Figure 4C), indicating a coordinated systemic inflammatory response, specifically activation of the innate immune system and hepatic acute-phase response.

**Figure 4.**
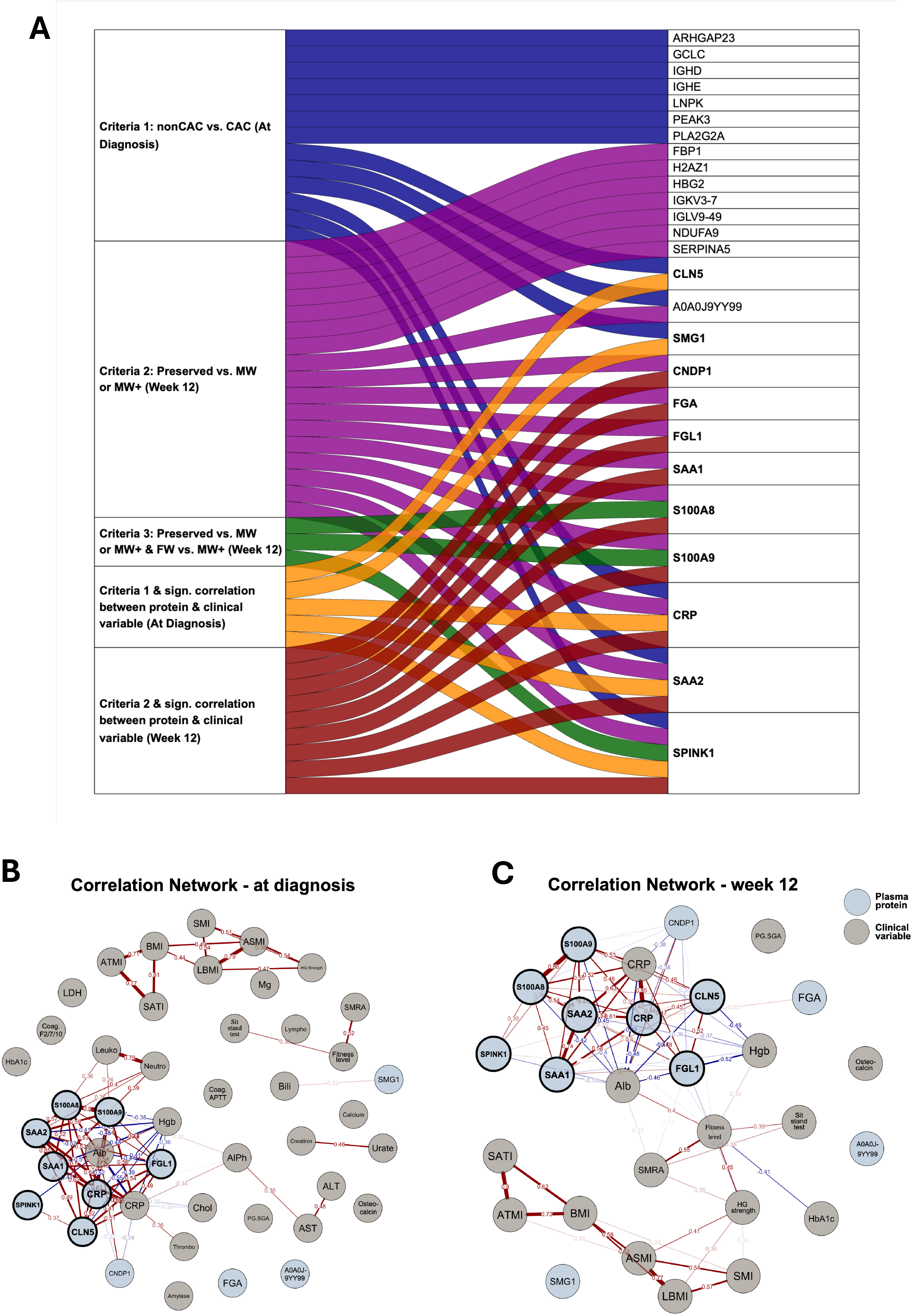
Target-selection workflow identifies potential drivers of NSCLC CAC and muscle wasting from the proteomic dataset. **A:** Three criteria formed the basis of the target selection (see Methods for details). Twenty-six proteins met at least one criterion; twelve met at least two criteria, and all but A0A0J9YY99 could be further analyzed for network clustering and clinical-variable association (bold). **B-C:** Correlation networks at diagnosis (B) and week 12 (C): proteins (blue) and clinical variables (grey) connected by lines showing positive (red) or negative (blue) Kendall’s tau correlations. Eight proteins met at least two criteria and showed network clustering with positive correlations at both timepoints (black circles, bold): CLN5, FGL1, S100A8, S100A9, SAA1, SAA2, SPINK1, and CRP. Sit Stand Test=30-second sit-to-stand test. HG Strength=hand grip strength (kg). BMI=body mass index. LBMI=lean body mass index. SMI=skeletal muscle index (cm²/m²). SMRA=skeletal muscle radiation attenuation. ASMI=appendicular skeletal muscle index. ATMI=adipose tissue mass index. SATI=subcutaneous adipose tissue index. Hgb=hemoglobin. Alb=albumin. HbA1c=glycated hemoglobin. CRP=C-reactive protein. Calcium=ionized calcium (free). Leuko=leukocytes. Thrombo=thrombocytes. Neutro=neutrophils. Lympho=lymphocytes. Bili=bilirubin. ALT=alanine aminotransferase. AlPh=alkaline phosphatase. Coag.F2/7/10=coagulation factors II, VII, X. AST=aspartate aminotransferase. LDH=lactate dehydrogenase. Coag.APTT=activated partial thromboplastin time. Chol=cholesterol. Mg=magnesium. PG-SGA=patient-generated subjective global assessment.

Together^45^, our comprehensive plasma proteomics analyses identified distinct molecular signatures of skeletal muscle wasting, specifically identifying S100A8, S100A9, CRP, FGL1, SPINK1, CLN5, SAA1 and SAA2 as potential regulators of muscle wasting across the cachexia trajectory in patients with advanced stage NSCLC.

### SAA1 and SAA2 decrease myotube width and contractile properties in human muscle

Having identified distinct molecular signatures of skeletal muscle wasting, we next functionally annotated and validated the roles of these circulating proteins in human muscle. To assess their pro-atrophic potential, we treated human myocytes with recombinant forms of the eight candidate proteins (1 µg/ml, per previously used doses^32^) and quantified myotube width during differentiation (Figure 5A). Of these (Figure S3A), recombinant SAA1 and SAA2 reduced myotube width by 41% and 47%, respectively, on day 4 versus vehicle-treated controls (Figure 5B; representative images in Figure 5C). In contrast, recombinant human S100A8/A9 (hybrid), CRP, FGL1, SPINK1, and CLN5 did not affect myotube width in vitro at the tested doses, while the positive control, recombinant TGF-β1^33^ reduced myotube width by 60% (Figure S3A).

**Figure 5.**
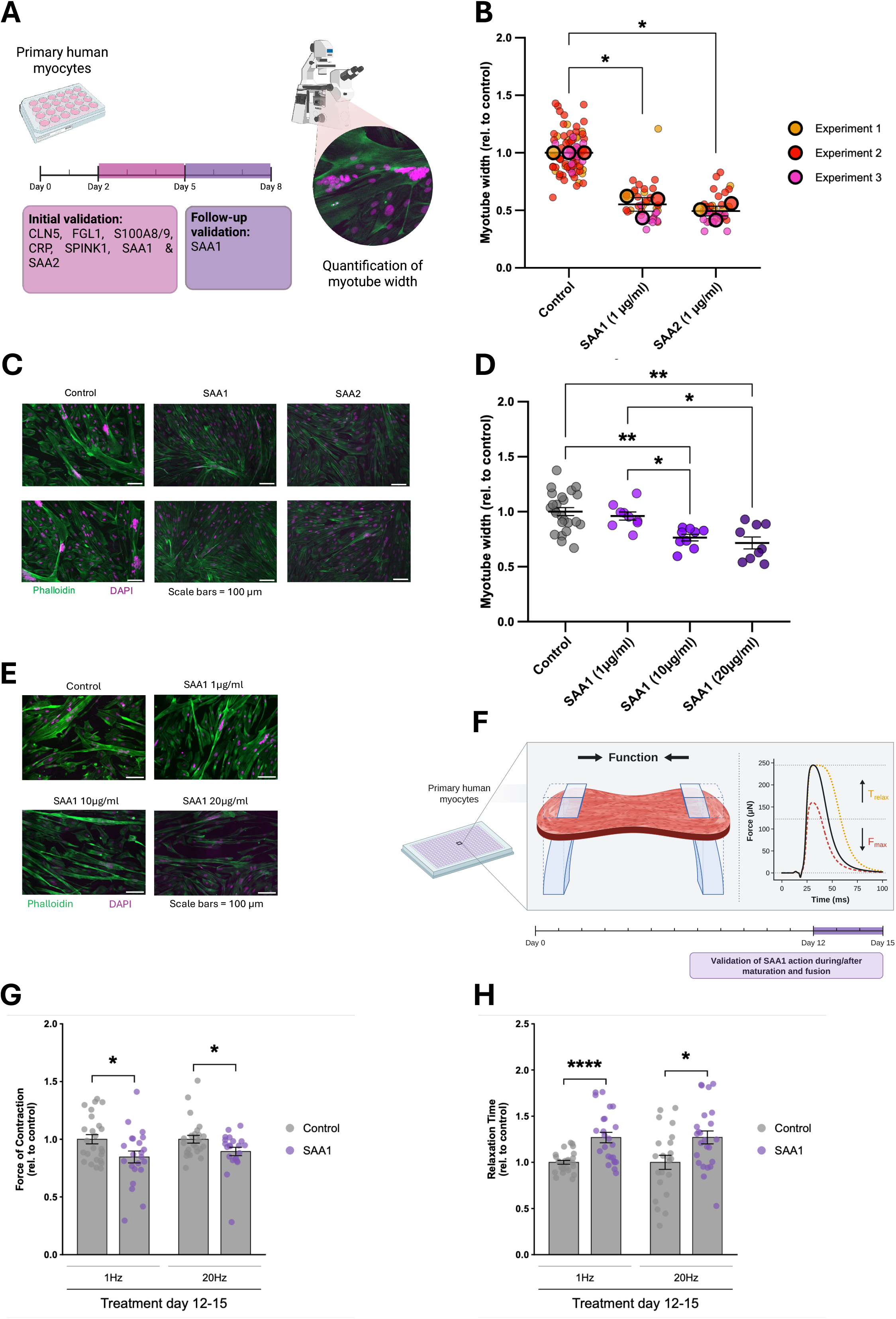
SAA1 and SAA2 impair human myotube development and contractile function. **A:** Schematic overview: primary human myocytes were treated during differentiation (day 2-5; initial screen: CLN5, FGL1, S100A8/9, CRP, SPINK1, SAA1, SAA2) or after maturation (day 5-8; follow-up: SAA1), followed by quantification of myotube width. **B:** Super-plot of pooled mean myotube width (relative to control) after treatment with BSA (control), recombinant SAA1, or SAA2 (1 µg/mL) on day 2-5. Large dots: biological replicates; small dots: technical replicates. Ratio-paired t-test of experiment means vs. control. **C:** Representative images of myotubes from (B). **D:** Dose-response validation in mature myotubes (day 5-8 treatment): width (relative to control) after SAA1 at 1, 10, or 20 µg/mL. Kruskal-Wallis test with Holm-adjusted Dunn’s multiple comparisons. **E:** Representative images (day 5-8 treatment). **F:** Schematic of the contractile-function assay: bioengineered 3D human myotubes were electrically stimulated and force of contraction measured (maximal force, Fmax; relaxation time, Trelax). SAA1 was given on day 12-15 (after maturation). **G:** Force of contraction (relative to control) at 1 Hz and 20 Hz after SAA1 (10 µg/mL) on day 12-15. Mann-Whitney test per day and frequency; significance levels shown. **H:** Relaxation time (relative to control) at 1 Hz and 20 Hz after SAA1 (10 µg/mL) on day 12-15. Mann-Whitney test per day and frequency. SAA1=serum amyloid A1. SAA2=serum amyloid A2. Magenta=DAPI (nucleus). Green=phalloidin (actin). Scale bars=25 µm. BSA=bovine serum albumin. Hz=hertz. µg=micrograms. mL=milliliters. *p<0.05, **p<0.01. ns=not significant.

We next investigated whether SAA also reduced myotube width in more mature myotubes, post differentiation and fusion. To this end, we quantified the effect of increasing SAA1 concentrations (0, 1, 10, 20 µg/ml) on myotube width in myotubes differentiated for 8 days. In these matured myotubes, we observed a dose-dependent 24–25% decrease in width (Figure 5D; representative images in Figure 5E). Having established SAA’s atrophic effect in mature human myotubes, we next assessed the functional consequences of SAA treatment using bioengineered 3D human muscle bundles^34^ (Figure 5F). SAA lowered contractile force at 1Hz by 15% in 3D muscle tissues (Figure 5G). SAA also slowed twitch relaxation time (1Hz), and tetanic relaxation time (20Hz) (Figure 5H). We observed no effect on activation time (Figure S3B). Representative curves are shown in Figure S3C,D. Together, these results indicate that SAA reduces myotube width and impairs twitch force production and relaxation time, consistent with preclinical cancer cachexia^35,36^. These detrimental effects are indicative of skeletal muscle dysfunction, key components of cachexic muscle^6^. These findings suggest that SAAs are potent drivers of human myotube atrophy and contractile function, providing functional annotation for their roles in regulating skeletal muscle mass.

### SAA secretion is driven by IL-6-receptor mediated signaling

Having established SAA1 and SAA2 as drivers of muscle wasting and associated with systemic inflammation, we next sought to delineate the upstream signaling events governing the CAC-associated increase in circulating SAA levels. In preclinical studies, cytokines, such as IL-1β, IL-6 and TNF-a are proposed as drivers of hepatic SAA production^37–39^. IL-6 correlates positively with weight loss in patients with NSCLC^40^ and negatively with survival in patients with pancreatic ductal adenocarcinoma (PDAC)^41^, raising the possibility that some of these systemic effects involve hepatic SAA production. To determine the individual and combined effects of these cytokines in SAA-release, we treated primary mouse hepatocytes with IL-6, IL-1β and TNF-a alone or a combination. Only IL-6 treatment alone elicited a 44% and 99% increase in SAA secretion for the low dose and high dose respectively, while the combination treatment had an additive effect, increasing SAA concentrations by 4-9 fold to 1µg/ml (Figure 6A), which associated with elevated levels of total STAT3 protein phosphorylated STAT3 (Figure 6B) in line with their role in cytokine signaling in hepatocytes. These results show that IL-6, and in particuliar in combination with other cytokines, evokes SAA secretion by primary hepatocytes. In agreement, liver tissues from mice with preclinical PDAC-associated cachexia displayed a time-dependent upregulation of *Stat3* and *Socs3*, with a shared temporal signature to *Saa1* and *Saa2* mRNA (Figure 6C and Figure S3E-G). These data support a model where cytokines like IL-6 drive STAT3-mediated SAA1/2 secretion, potentially restrained by SOCS3 negative feedback^42^. This aligns with the IL-6/STAT3 axis proposed to drive cachexia across multiple cancers^5,43^.

**Figure 6.**
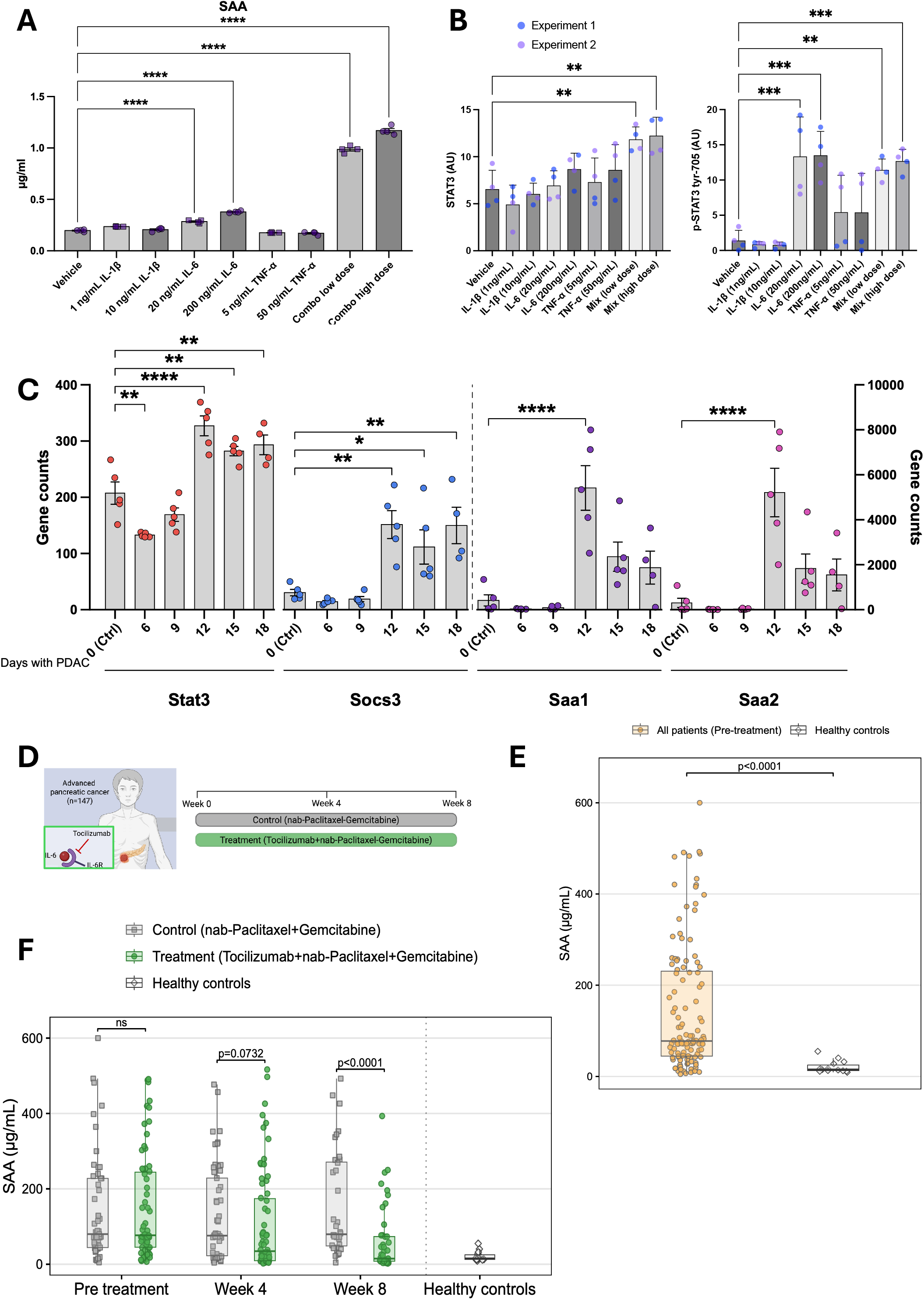
Cytokine-driven hepatocyte SAA production, and SAA in patients with pancreatic cancer with or without tocilizumab (IL-6 blockade) **A:** SAA levels (ELISA, µg/mL) in conditioned medium from primary mouse hepatocytes after stimulation with IL-1β (1, 10 ng/mL), IL-6 (20, 200 ng/mL), TNF-α (5, 50 ng/mL), or low-/high-dose combinations, vs. vehicle (n=4/group). Bars: mean+SEM; dots: individual replicates. One-way ANOVA with Dunnett’s test vs. vehicle. **B:** Western blot of hepatocyte phospho-STAT3 and SOCS3. **C:** Hepatic Stat3, Socs3, Saa1, and Saa2 gene counts (RNA-seq) in sham (Ctrl) and PDAC mice at days 6, 9, 12, 15, and 18 with PDAC (n=4-5 mice/timepoint. Normalized counts are expressed as CPM (Counts per million). Bars: mean+SEM; dots: individual mice. One-way ANOVA with Dunnett’s test vs. day 0, per gene. **D:** Trial schematic: 146 patients with pancreatic cancer (PC) randomized to standard treatment (nab-paclitaxel+gemcitabine) or standard treatment plus 8 weeks of tocilizumab; blood drawn at week 0, 4, and 8. **E:** SAA levels (ELISA) in PC patients pre-treatment vs. healthy controls. Wilcoxon rank-sum test; p value shown. **F:** SAA levels in PC patients with and without tocilizumab, and healthy controls, at pre-treatment, week 4, and week 8. Wilcoxon rank-sum tests; p values shown. CAC=cachexia. PC=pancreatic cancer. PDAC=pancreatic ductal adenocarcinoma. KPC=murine PDAC model. IL-6=interleukin-6. IL-1β=interleukin-1β. TNF-α=tumor necrosis factor-α. SAA=serum amyloid A. CM=conditioned medium. µg=micrograms. mL=milliliters. ng=nanograms. Ctrl=control. SEM=standard error of the mean. *p<0.05, **p<0.01, ****p<0.0001.

To determine whether this IL-6-dependent SAA regulatory axis operates as a conserved mechanism across distinct cachexia-prone malignancies, we sought independent, in vivo pharmacological validation. We evaluated SAA levels in a separate clinical cohort of patients with locally advanced or metastatic PC receiving first-line nab-paclitaxel and gemcitabine, either with or without the humanized anti-IL-6 receptor antibody tocilizumab (Table 2, Figure 6D)^44^. We found that patients with PC had markedly (5.9-fold) elevated SAA levels compared to an age-matched healthy control group (Figure 6E). Notably, SAA levels trended 18% lower in the tocilizumab group versus nab-paclitaxel/gemcitabine alone at 4 weeks (p=0.0732), reaching a significant 66% reduction by week 8 (Figure 6F). Accordingly, there was an 81% decrease in the tocilizumab-treated group (week 8 vs. pre-treatment, p<0.001). Thus, IL-6 receptor blockade normalized total plasma SAA in patients with PC (Figure S4A) and was accompanied by attenuated muscle mass loss and improved survival in the tocilizumab-treated group^18^, supporting an IL-6–SAA axis in CAC. We also correlated circulating SAA levels with circulating IL-6 (Figure S4B) and CRP (Figure S4C) and found positive associations between SAA and both of these inflammatory cytokines, similar to what we observed for CRP in patients with NSCLC (Figures S4D).

**Table 2.** Characteristics of patients with pancreatic cancer (randomized to control or tocilizumab) and healthy controls. Data are n (%) or mean (SD), unless otherwise noted; percentages use the number of patients in each row as the denominator. Control=nab-paclitaxel+gemcitabine. Treatment=tocilizumab+nab-paclitaxel+gemcitabine. PS=Eastern Cooperative Oncology Group Performance Status. BMI=body mass index. CRP=C-reactive protein. IL-6=interleukin-6.

| **Variable** | **Control**<br>**(nab-**<br>**Paclitaxel+Gemcitabine)** | **Treatment**<br>**(Tocilizumab+nab-**<br>**Paclitaxel+Gemcitabine)** | **Healthy**<br>**Controls** |
| --- | --- | --- | --- |
| **Biological sex** | N = 70 | N = 76 | N = 14 |
| Male, N (%) | 42 (60) | 43 (56.6) | 8 (47.1) |
| Female, N (%) | 28 (40) | 33 (43.4) | 6 (52.9) |
| **Age at**<br>**randomisation** |  |  |  |
| Years,<br>median (min-max) | N = 70<br>67 (36-84) | N = 76<br>68 (34-84) | N = 14<br>69 (56-78) |
| **Cachexia status**<br>**at time of**<br>**diagnosis** | N = 67 | N = 72 |  |
| $\geq 5\%$, N (%) | 49 (73.13) | 45 (62.5) | |
| $< 5\%$, N (%) | 18 (26.87) | 27 (37.5) | |
| **PS** | N = 70 | N = 74 |  |
| 0, N (%) | 44 (62.86) | 48 (63.16) |  |
| 1, N (%) | 26 (37.14) | 36 (36.84) |  |
| 2, N (%) | 0 (0) | 0 (0) |  |
| **BMI** | N = 70 | N = 76 | N = 14 |
| Mean (SD) | 25.12 (4.30) | 24.84 (4.02) | 25.23 (6.47) |
| **CRP (mg/L)** | N = 70 | N = 76 | N = 14 |
| Median (min-max) | 25 (3-244) | 31 (3-259) | 1 (0-9) |
| **Albumin (g/L)** | N = 67 | N = 75 |  |
| Median (min-max) | 39 (19-49) | 37 (26-46) |  |
| **IL-6 (pg/ml)** | N = 68 | N = 73 |  |
| Median (min-max) | 7.6 (1.4-9208) | 8.75 (10.4-103.1) |  |

Together these preclinical and clinical data indicate that SAA secretion is driven by IL-6-receptor mediated signaling in CAC.

### Baseline SAA levels predict functional decline and poor survival in patients with lung cancer and pancreatic cancer

Having established an IL-6–associated SAA axis linked to muscle wasting, we next investigated whether SAA levels would clinically relate to declined muscle function and survival in patients with advanced lung and pancreatic cancer.

First, we analyzed our cohort of patients with NSCLC. SAA1 and SAA2 are highly homologous, and their plasma abundances were closely correlated (Figure S5A). Consistent with the pro-atrophic effects of recombinant SAA observed in human myotubes, SAA levels at diagnosis measured by ELISA were elevated (2.9-fold) in the MW+ compared to Preserved patients (Figure 7A) corroborated by the proteomics data showing increase of both SAA1 (Figure 7B) and SAA2 (Figure 7C) at diagnosis in the MW+ compared to Preserved NSCLC patients. Additionally, when stratifying the patients based on severity of muscle wasting, the patients with the most muscle wasting also had the highest SAA levels at diagnosis compared to the patients with the least muscle wasting (Figure 7D). Furthermore, rising SAA1 tracked inversely with sit-to-stand performance (Figure S5B). Positioning SAAs at the nexus of physical decline and systemic inflammation, diagnostic SAA1 and SAA2 correlated inversely with hemoglobin (bone marrow suppression; Figure S5C) and albumin (metabolic/nutritional disturbance; Figure S5D), and positively with CRP (inflammation) and neutrophils (immune response) (Figures S5E). Longitudinally, increases in both SAAs consistently associated with declining albumin (Figure S5F) and elevated CRP (Figure S5G).

**Figure 7.**
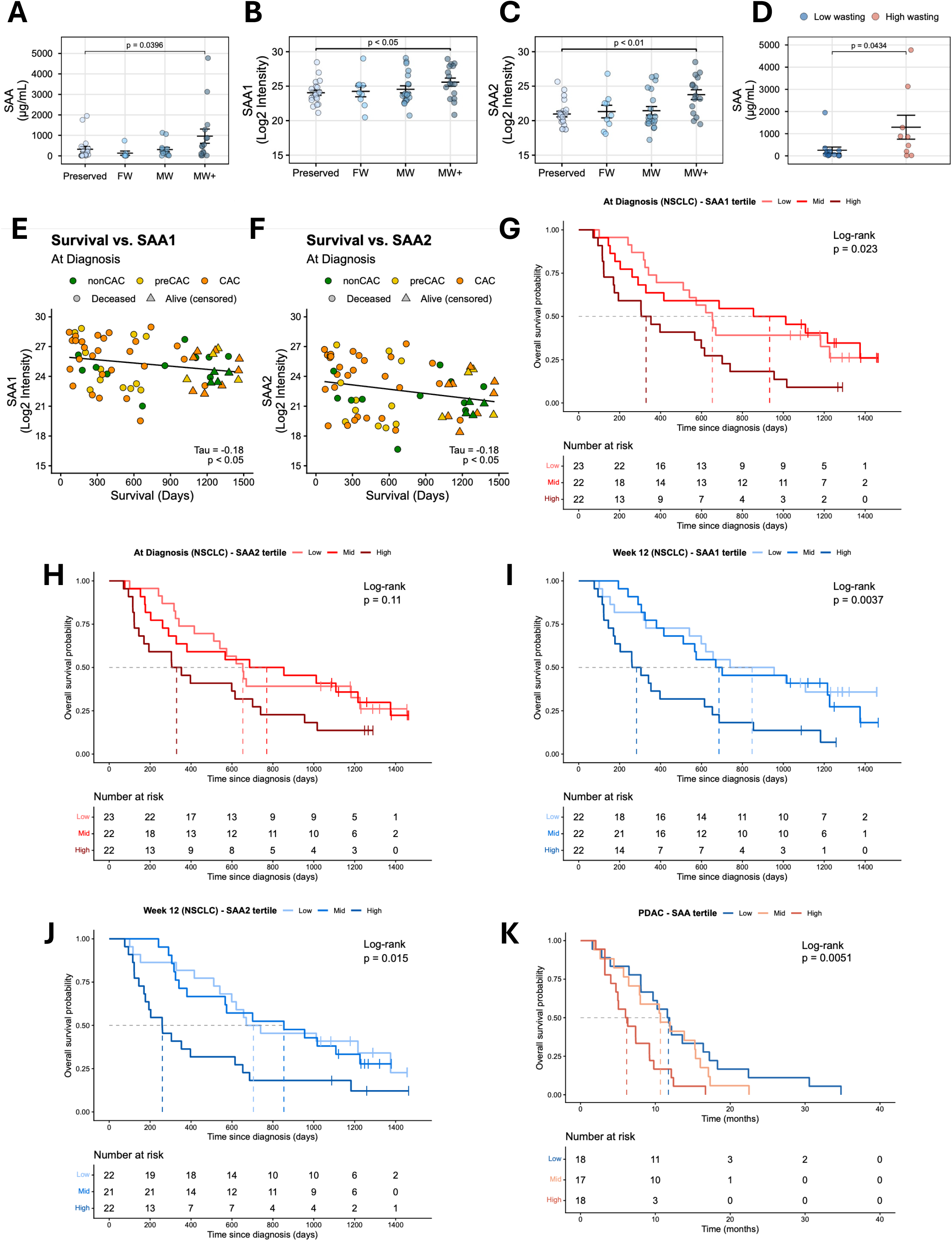
Circulating SAA associates with wasting phenotype and overall survival. **A:** Plasma SAA (µg/mL) at diagnosis by week-12 wasting phenotype. Dots: individual patients; lines: mean±SEM. Wilcoxon rank-sum test, Preserved vs. MW+. **B-C:** MS-measured SAA1 (B) and SAA2 (C) levels (log2 intensity) at week 12, by wasting phenotype. **D:** Plasma SAA (ELISA) at diagnosis in patients grouped by low or high wasting (week-12 delta-muscle-area tertiles; middle tertile excluded). Wilcoxon rank-sum test. **E-F:** Correlation of overall survival (days from diagnosis) with diagnostic SAA1 (E) and SAA2 (F) (log2 intensity); n=67: deceased (circles, n=51), alive/censored (triangles, n=16); colored by CAC status. **G-H:** Kaplan-Meier overall survival by diagnostic SAA1 (G) and SAA2 (H) tertile. Log-rank test. **I-J:** Kaplan-Meier overall survival by week-12 SAA1 (I) and SAA2 (J) tertile. Log-rank test. **K:** Kaplan-Meier overall survival by pre-treatment SAA (ELISA) tertile in PC patients (control group). Log-rank test. Numbers at risk are shown below each plot (G-K). SAA=serum amyloid A. SAA1=serum amyloid A1. SAA2=serum amyloid A2. Log2 intensity=log2-transformed concentration. nonCAC=no cachexia. preCAC=pre-cachexia. CAC=cachexia. Preserved=no wasting. FW=fat wasting. MW=muscle wasting without weight loss. MW+=muscle wasting with weight loss. MS=mass spectrometry. NSCLC=non-small cell lung cancer. Tau=Kendall’s tau, a non-parametric rank-correlation coefficient. µg=micrograms. mL=milliliters.

Having established SAA1 and SAA2 as clinical markers of muscle wasting in NSCLC, we next asked whether plasma levels of SAA1 (Figure 7E) and SAA2 (Figure 7F) correlate with overall survival. At diagnosis, both isoforms correlated inversely with survival (Kendall’s tau = −0.18; both p < 0.05), and Kaplan-Meier analysis confirmed that elevated SAA1 (Figure 7G) and SAA2 (Figure 7H) predict poorer overall survival in NSCLC. This prognostic value strengthened by week 12: patients in the highest SAA1 tertile had a median survival of only 283 days, versus 701 (mid) and 740 (low) days (log-rank p = 0.0037; Figure 7I), and one- and two-year survival fell to 36% and 18% in the highest tertile, versus 73% and 55% in the lowest, with similar direction for SAA2 (Figure 7J).

We then tested whether SAA levels similarly predicted survival in our validation PC cohort. High diagnostic SAA (measured by ELISA, which does not distinguish SAA1 from SAA2) predicted markedly reduced survival in PC as well (Figure 7K), mirroring trends for elevated IL-6 (Figure S5H) and CRP (Figure S5I). Together, these clinical data indicate that in cachexia-prone cancers, elevated circulating SAA correlates with accelerated muscle wasting during NSCLC treatment, functional decline and poorer survival in both NSCLC and PC.

## DISCUSSION

By integrating longitudinal clinical phenotyping, plasma proteomics, functional studies in human myotubes, and a pharmacodynamic analysis of IL-6 receptor blockade, we identify SAA1 and SAA2 as acute-phase proteins linked to muscle wasting and survival in human cancer. Plasma proteomic signatures at diagnosis predicted distinct wasting trajectories in NSCLC, and SAA1 and SAA2 directly modulated human myotube mass and function. Elevated SAA associated with poor survival in both the NSCLC and PC cohorts, and in PC, tocilizumab treatment reduced circulating SAA, suggesting that SAA is at least partly regulated by IL-6 receptor signaling. Together, these findings support an IL-6-SAA-muscle wasting axis that warrants further mechanistic and interventional validation.

While cross-sectional findings underscore the clinical burden of cachexia ^45–51^ and the urgent need for a deeper biological understanding of its pathogenesis, the characterization of cachexia has historically been limited by a reliance on weight loss alone^31,52^. This dependency often obscures the specific loss of skeletal muscle mass. We demonstrate that dynamic wasting trajectories provide a more precise molecular map of disease progression, capturing proteomic shifts overlooked by single-point assessments. Integrating concordant longitudinal CT and DXA imaging enabled us to define distinct signatures for cachexia at diagnosis and specific phenotypic clusters of fat and muscle wasting. These findings reveal that clinical heterogeneity in wasting is associated with divergent, tissue-specific molecular programs, previously uncharacterized at this resolution. At diagnosis, a systemic pro-inflammatory shift defines the cachectic proteome, reinforcing the classification of cachexia as a systemic inflammatory disease.

Through a clinically anchored selection workflow, we identified eight proteins upregulated in patients with skeletal muscle loss: S100A8, S100A9, CRP, FGL1, SPINK1, CLN5, SAA1, and SAA2. S100A8/A9, CRP, and FGL1 have previously been linked to cancer-associated wasting or prognosis ^53–58^, whereas SPINK1 and CLN5 have been studied mainly in tumor biology ^59–62^ and have no reported role in cachexia. SAA1 and SAA2 are hepatic acute-phase proteins. SAA1 has been associated with murine cancer cachexia and reduces myotube width in murine muscle cells^32,43,63^, and in humans with muscle atrophy in non-alcoholic fatty liver disease ^64^ and poor outcome in NSCLC^65^. SAA2 is a proposed lung cancer plasma marker^66^, but its association with muscle atrophy or cachexia has, to our knowledge, not been reported.

SAA1 and SAA2 signal through Toll-like receptors (TLRs) in murine models^67,68^, and preliminary data suggest a similar mechanism in human muscle^69^. We extend these findings by showing that SAA1 and SAA2 directly induce atrophy and contractile dysfunction in human myotubes. These data points to impaired excitation-contraction coupling, with reduced Ca²⁺ release from the sarcoplasmic reticulum per stimulus and slower Ca²⁺ reuptake, rather than a defect in the contractile apparatus. It is consistent with the altered Ca²⁺ handling we observed in skeletal muscle from patients with NSCLC^70^ and with the oxidized, Ca²⁺-leaky RyR1 channels suggested to drive cancer- and aging-associated muscle weakness independently of atrophy^71,72^. Whether SAA acts through TLRs and Ca²⁺-handling proteins in human muscle remains to be tested.

To validate the regulatory hierarchy of this axis in vivo, we leveraged an independent pharmacological intervention cohort of patients with advanced PC. The profound suppression of SAA levels following IL-6 receptor blockade with tocilizumab not only confirms the systemic IL-6-SAA axis but crucially demonstrates that this inflammatory-metabolic relay is a conserved mechanism across multiple cachexia-prone tumor types. In this paradigm, generalized tumor-driven IL-6 signaling triggers hepatic SAA secretion, which subsequently operates as a direct effector of peripheral muscle catabolism. Collectively, our findings from longitudinal plasma proteomics, functional myotube assays, independent IL-6 receptor blockade analysis, and clinical evidence linking elevated SAA to reduced survival, underscore the IL-6-SAA axis as a compelling pathway of intervention and a potential critical determinant of host frailty.

While this study provides a comprehensive prospective dataset, several limitations warrant consideration. The study included different cohorts of patients with NSCLC and PC with different experimental and clinical contexts. The proposed IL-6-SAA-muscle wasting axis should be interpreted as a biologically supported model rather than a fully proven causal pathway in patients with cancer. Despite low loss to follow-up, our cohort may be subject to selection bias, as only participants with the energy to adhere to the protocol were included, likely underrepresenting patients with higher disease burden or lower performance status^73^.

Strengths of this study include its longitudinal design, which enabled the characterization of intra-patient changes through extensive clinical phenotyping and dual-modality imaging (CT and DXA) integrated with high-resolution systemic molecular signatures of cachexia. By utilizing clinical CT-derived wasting phenotypes, we established a powerful framework for identifying plasma markers specific to muscle loss. Our cohort of newly diagnosed, advanced-stage NSCLC and PC patients from large cancer centers with universal health coverage suggests that our populations are representative of a broader background population. We minimized experimental variability by employing standardized assessment personnel and centralized facility analysis for all samples. Furthermore, temporal monitoring of nutritional risk every three weeks allowed for individualized support, mitigating treatment-related symptoms that could otherwise confound the wasting condition.

In summary, we define the molecular heterogeneity of cancer cachexia by linking distinct adipose and muscle wasting phenotypes to specific plasma proteomic signatures. Crucially, we identify SAA1 and SAA2 as IL-6-driven mediators of this process that directly induce human myotube atrophy and dysfunction, establishing a functional link between systemic inflammation and skeletal muscle catabolism. In vivo pharmacological blockade with the IL-6 receptor antagonist tocilizumab validates this circuit, confirming that hepatic SAA production is a direct downstream effector of IL-6 signaling. Ultimately, our findings elevate the IL-6– SAA1/2 axis from a correlative biomarker signature to a functionally targetable driver of cancer cachexia. Disrupting this inflammatory-metabolic relay offers a promising, precision-medicine strategy to preserve muscle mass, mitigate treatment-limiting toxicity, and extend survival in patients with advanced malignancies^74^.

## Acknowledgments

We thank research assistant Nanna Hahn for data collection and patient contact throughout the study, and Nanna Hahn, Nicoline Resen Andersen and Simon Bech Petersen for handling of biological samples. We thank nurses Marie Aller and Fie Sylvest and physiotherapist Magnus Nygaard Bech for managing patient flow and support, and physiotherapists Malthe Mollerup-Degn Hermann, Andreas Ubberup Larsen and Barbara Bordignon for supporting patients during physical assessments. We thank all nurses and medical doctors at the Department of Oncology, Copenhagen University Hospital – Rigshospitalet, for supporting inclusion and the participating patients and relatives; senior radiology consultant Thomas Skårup Kristensen for guidance on radiology data acquisition; assistant professor Wouter van de Worp for guidance on data handling and interpretation; and research assistant Ralph Brecheisen for body composition analysis. Mass spectrometry-based proteomics was performed by the Proteomics Research Infrastructure (PRI), University of Copenhagen, supported by the Novo Nordisk Foundation (NNF19SA0059305). We thank the MyoLine platform of the Center of Research in Myology and the Myobank-AFM biobank of the Institute of Myology. Above all, we are deeply grateful to the participating patients and relatives who made this study possible.

## AUTHORS CONTRIBUTIONS

Conceptualization: J.S. and L.S. Data curation: J.S., J.M., B.C, J. C., C.K., Z.O., C.V., A.L., T.C., R.M., E.B. and A.H. Formal analysis: J.S., J.M., C.K., Z.O., A.L., T.C., R.M., C.L, and L.S. Funding acquisition: J.S., R.M and L.S. Investigation: J.S., Z.O., I.C., E.A.R, J.S.J., and E.B. Methodology: J.S., E.B., I.C., J.S.J, Z.O., and L.S. Project administration: J.S. and L.S. Resources: L.S. and S.L. Software: J.S., C.V., E.B. and L.S. Supervision: K.S, T.Z, L.S. Validation: J.S. Visualization: J.S., C.V., Z.O., and E.B. Writing original draft: J.S., Z.O., L.S., Writing review & editing: all authors.

---

**Figure S1.**
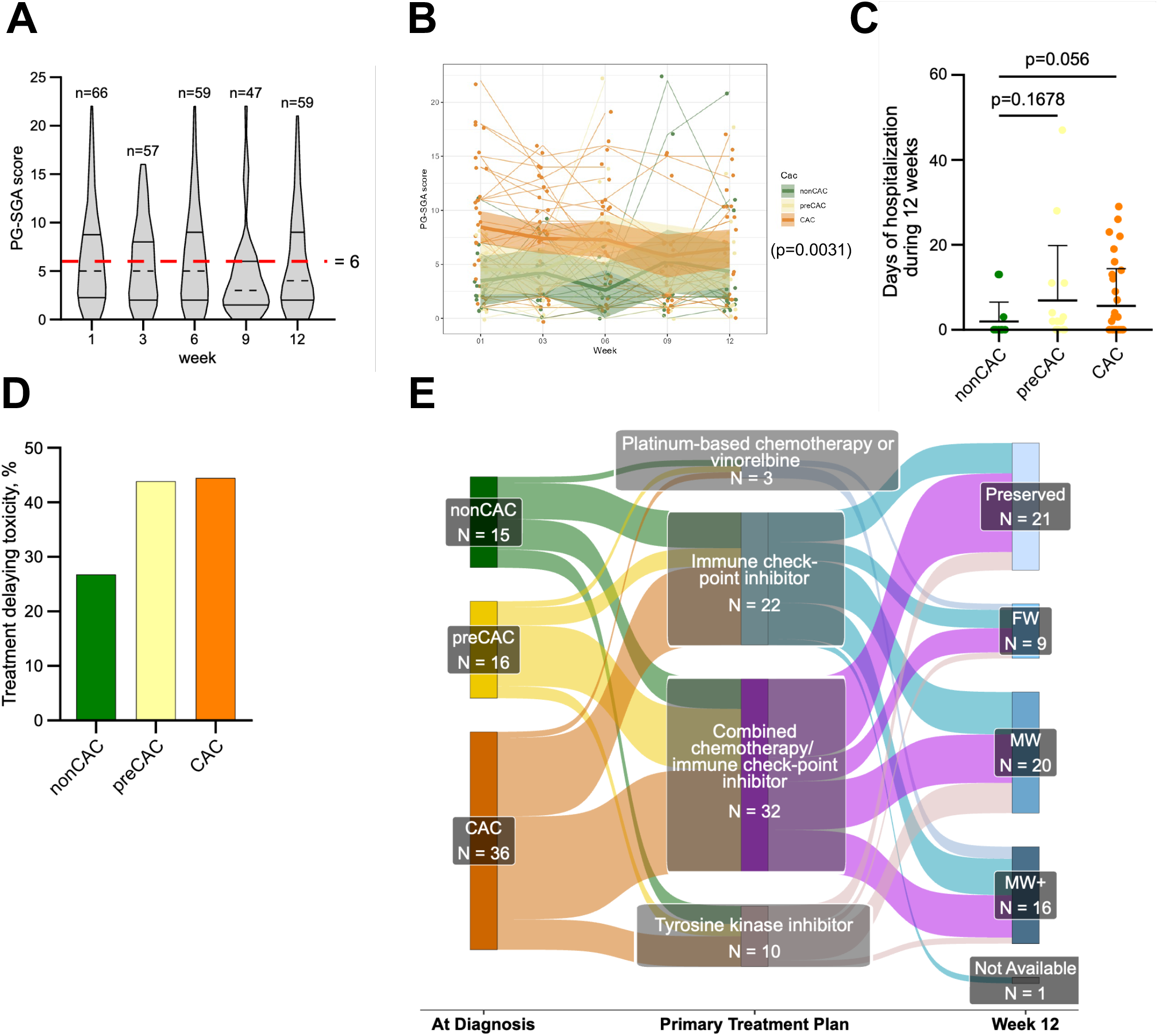
Nutritional risk, hospitalization, and treatment-delaying toxicity by CAC group and wasting trajectory. **A:** PG-SGA scores at weeks 1, 3, 6, 9, and 12 (violin plots) with means (dark dotted lines), 95% CI (dark solid lines), and the nutritional-risk cutoff of 6 (red dotted line). Dunnett’s test compared week 1 to weeks 3, 6, 9, and 12 (47-66 patients per timepoint; 70-98% response rate). No significant differences were found. **B:** Individual PG-SGA trajectories by CAC status (nonCAC, green; preCAC, yellow; CAC, orange) across weeks 1-12. A mixed model estimated the effect of CAC status on PG-SGA scores during treatment (p=0.0031). PG-SGA=patient-generated subjective global assessment. nonCAC=no cachexia. preCAC=pre-cachexia. CAC=cachexia. **C:** Days of hospitalization over 12 weeks by CAC status at diagnosis (dots: individual patients; solid line: mean). Mean days: nonCAC 1.9, preCAC 6.9, CAC 5.6. Welch’s t-test: nonCAC vs. preCAC p=0.168; nonCAC vs. CAC p=0.056. **D:** Percentage of patients with treatment-delaying toxicity. For all patients during their 12 weeks of first-line treatment, it was recorded whether the oncologist responsible for the treatment postponed the next scheduled treatment cycle due to treatment-related toxicity. Patients who had their treatment postponed at least once for this reason were grouped as treatment-related toxicity.: nonCAC 26.7%, preCAC 43.8%, CAC 44.4%. Fisher’s exact test, nonCAC vs. preCAC+CAC, p=0.251. nonCAC=no cachexia. preCAC=pre-cachexia. CAC=cachexia. **E:** Sankey diagram showing CAC status at diagnosis, treatment group, and contribution to three wasting-phenotype trajectories over 12 weeks. Band width is proportional to patient number, so the widest bands represent the dominant clinical phenotypes. Treatment groups: platinum-based chemotherapy/vinorelbine, immune checkpoint inhibitor, combination chemotherapy/checkpoint inhibitor, or tyrosine kinase inhibitor. Wasting phenotypes were defined by CT-derived muscle and weight loss: Preserved (no wasting; <2.5% loss of both SMI and SATI), fat wasting (FW), muscle wasting without weight loss (MW), and muscle wasting with weight loss (MW+; total body weight loss >2.5% distinguishes MW from MW+). One patient was lost to follow-up CT at week 12; one patient with a technical CT error was classified as Preserved based on weight gain and unchanged DXA measures. CT=computed tomography. SATI=subcutaneous adipose tissue index. SMI=skeletal muscle index. WL=weight loss. BMI=body mass index. nonCAC=no cachexia. preCAC=pre-cachexia. CAC=cachexia.

**Figure S2.**
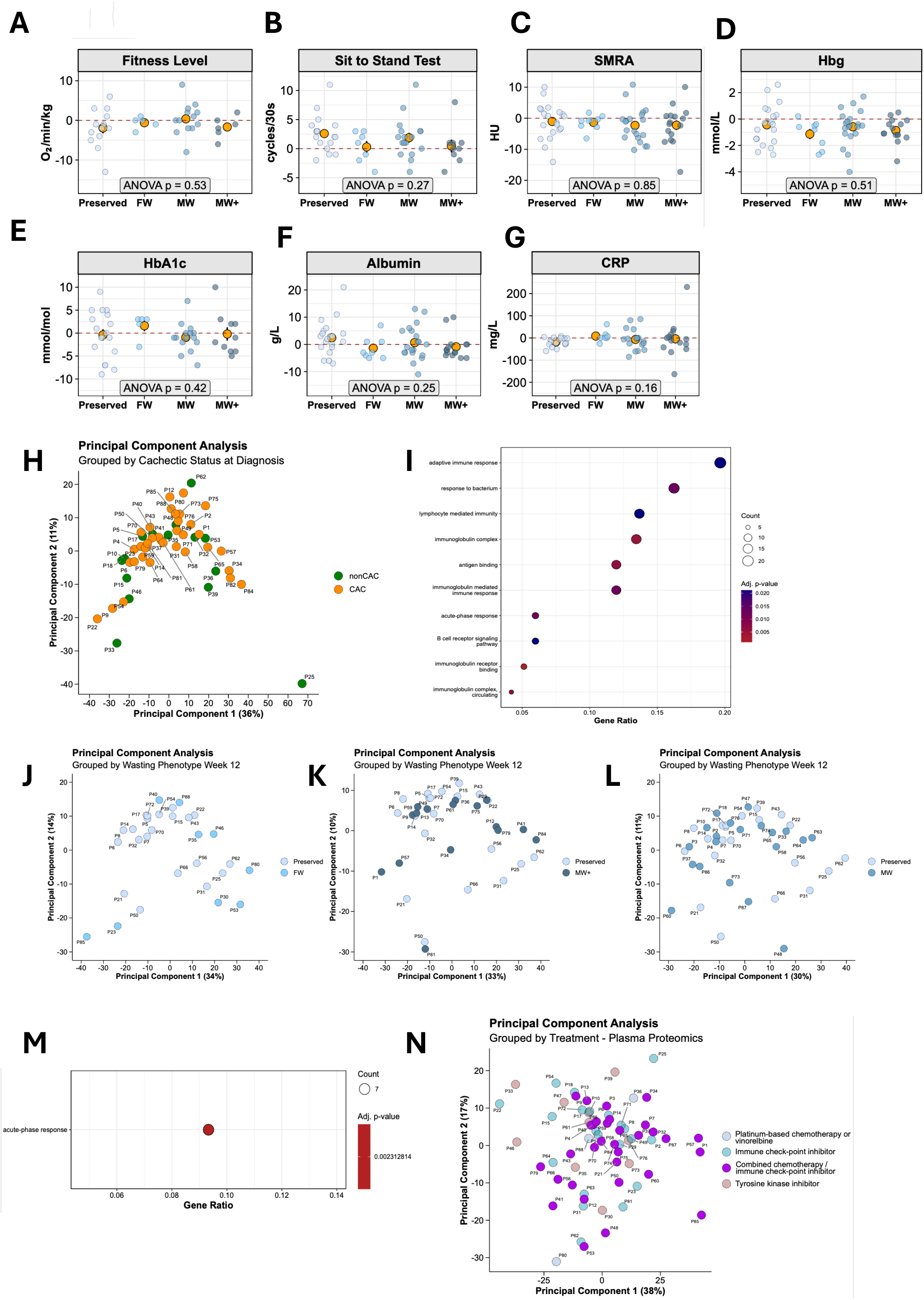
Principal component and over-representation analyses of the plasma proteome. **A-G:** Changes in clinical variables from diagnosis to week 12 by wasting phenotype: fitness level (A), sit-to-stand test (B), SMRA (C), hemoglobin (D), HbA1c (E), albumin (F), and CRP (G). Dots: individual patients (semi-transparent) and mean (orange); thin vertical lines: SEM. None of the variables differed significantly across phenotypes (one-way ANOVA). **H:** PCA of the diagnostic plasma proteome by cachexia status (nonCAC vs. CAC). **I:** Over-representation analysis of proteins upregulated in CAC at diagnosis. Color: significance; circle size: number of proteins; x-axis: gene ratio; y-axis: pathway. **J-L:** PCA of the week-12 plasma proteome: Preserved vs. FW (J), vs. MW (K), vs. MW+ (L). **M:** Over-representation analysis of proteins upregulated in MW+ vs. Preserved at week 12 (as in I). **N:** PCA of proteomic change (diagnosis to week 12) for all subjects, by treatment group. nonCAC=no cachexia. CAC=cachexia. Preserved=no wasting. FW=fat wasting. MW=muscle wasting without weight loss. MW+=muscle wasting with weight loss. SMRA=skeletal muscle radiation attenuation. HU=Hounsfield units. HbA1c=glycated hemoglobin. CRP=C-reactive protein. SEM=standard error of the mean.

**Figure S3.**
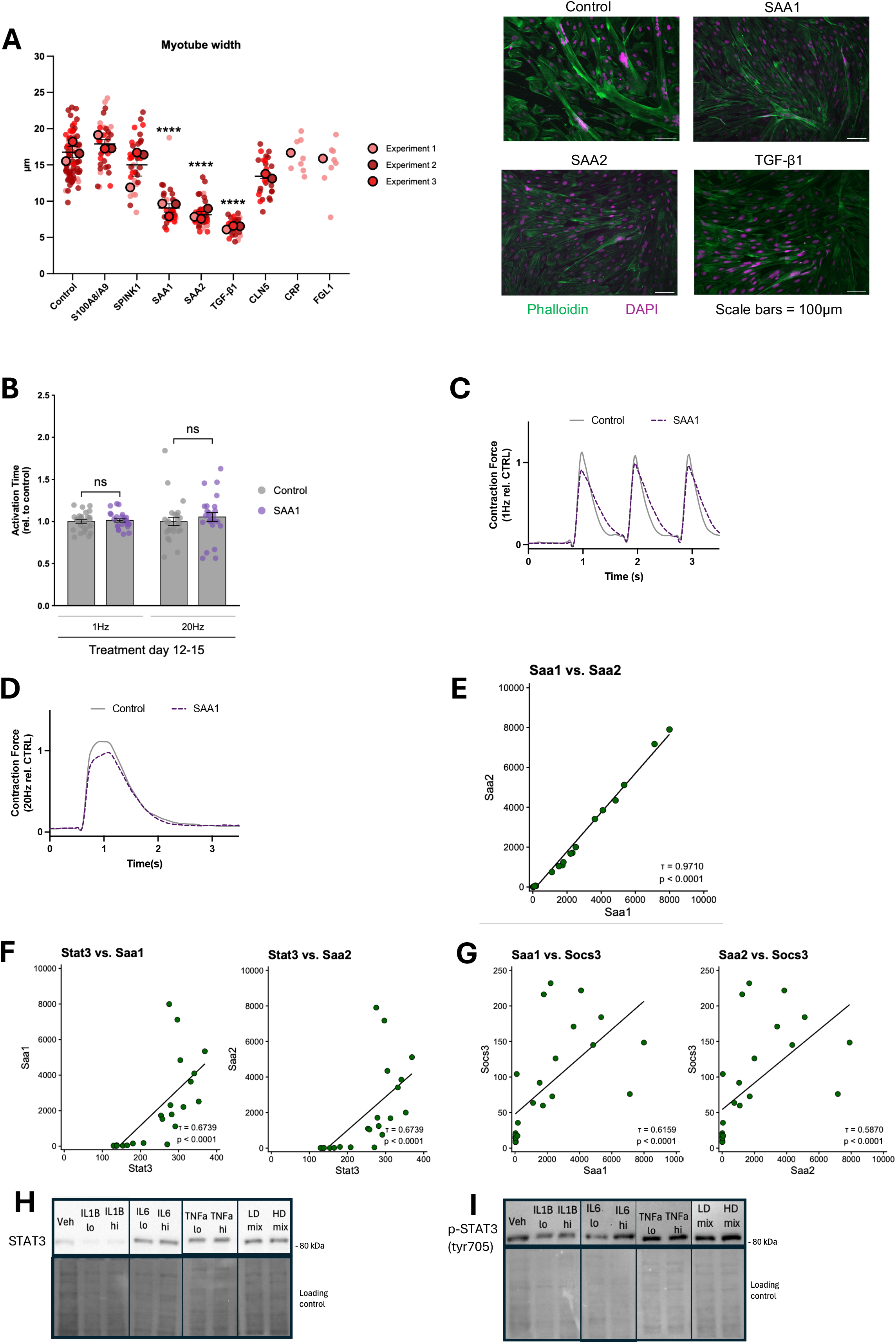
Myotube width across candidate proteins and contractile properties of SAA1-treated myotubes. **A:** Super-plot of pooled mean myotube width (µm) from myocytes treated on day 2-5 with BSA (control), S100A8/A9, SPINK1, SAA1, SAA2, TGF-β1, CLN5, CRP, or FGL1 (n=3; large dots: biological replicates, small dots: technical replicates). One-way ANOVA with Dunnett’s test vs. control; significant for SAA1, SAA2, and TGF-β1 (****p<0.0001). Right: representative images of myotubes treated with BSA (control), SAA1, SAA2, and TGF-β1. Green=phalloidin (actin). Magenta=DAPI (nucleus). Scale bars=25 µm. **B:** Activation time (relative to control) after electrically induced contraction at 1 Hz and 20 Hz in engineered human 3D muscle tissue treated with SAA1 (10 µg/mL) on day 12-15. Mann-Whitney test per day and frequency; all comparisons non-significant. **C-D:** Representative force traces at 1 Hz (C) and 20 Hz (D) for control and SAA1. BSA=bovine serum albumin. S100A8=S100 calcium-binding protein A8. S100A9=S100 calcium-binding protein A9. SPINK1=serine protease inhibitor Kazal type 1. SAA1=serum amyloid A1. SAA2=serum amyloid A2. TGF-β1=transforming growth factor beta 1. CLN5=ceroid-lipofuscinosis neuronal protein 5. CRP=C-reactive protein. FGL1=fibrinogen-like protein 1. Hz=hertz. ns=not significant. µm=micrometers. **E-G:** Correlations of hepatic gene counts (RNA-seq) in PDAC mice (day 6-18; n=24 samples): Saa1 vs. Saa2 (E), Stat3 vs. Saa1/2 (F), Socs3 vs. Saa1/2 (G). Kendall’s tau and p values shown; lines are linear-regression fits. PDAC=pancreatic ductal adenocarcinoma. **H-I:** Representative western blot image from hepatocytes treated with cytokines. Veh = Vehicle. IL1B lo= IL-1β (1ng/ml). IL1B hi= IL-1β (10ng/ml). IL6 lo=IL-6 (20 ng/ml). IL6 hi=IL-6 (200 ng/ml). TNFa lo = TNF-α (5ng/ml). TNFa hi = TNF-α (50ng/ml). LD mix=low dose mix (all cytokines at lowest stated dose). HD mix=high dose mix (all cytokines at highest stated dose).

**Figure S4.**
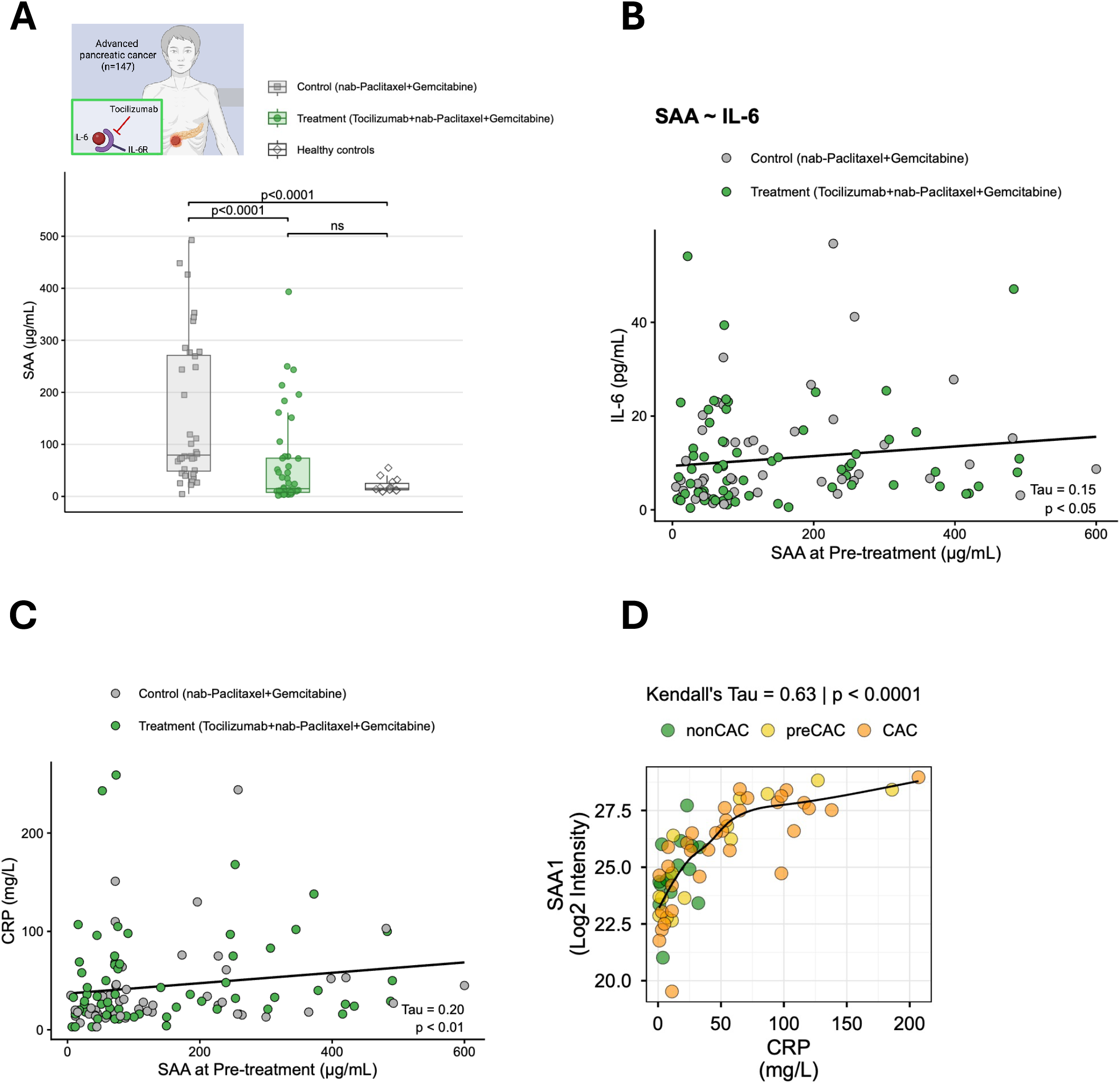
SAA, IL-6, and CRP in patients with pancreatic cancer with or without tocilizumab, and the IL-6/SAA axis. **A:** SAA levels at week 8 in control and treatment patients with pancreatic cancer, and healthy controls. Pairwise Wilcoxon rank-sum tests (Benjamini-Hochberg adjusted); p values shown. **B:** Correlation of SAA (ELISA) and IL-6 in patients with pancreatic cancer, pre-treatment. Kendall’s tau and p value shown. **C:** Correlation of SAA (ELISA) and CRP in patients, pre-treatment. Kendall’s tau and p value shown. **D:** Correlation of SAA1 (MS) and CRP in patients, pre-treatment. Kendall’s tau and p value shown.

**Figure S5.**
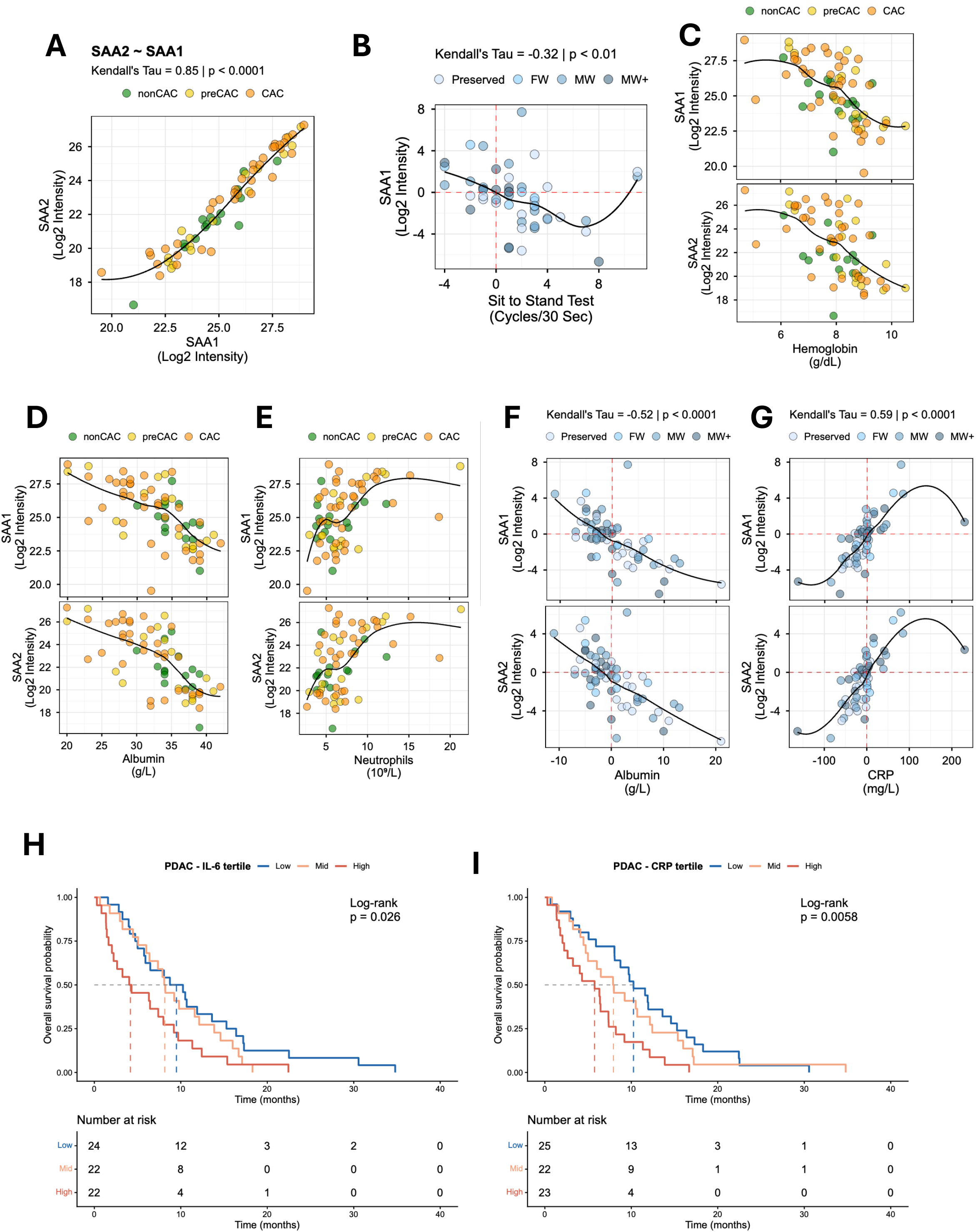
SAA1 and SAA2 in patients with NSCLC across CAC and wasting-phenotype groups, and survival in pancreatic cancer. **A:** Correlation of SAA1 and SAA2 plasma levels (MS) at NSCLC diagnosis, colored by CAC status. Kendall’s tau and p value shown. **B:** Correlation of change (diagnosis to week 12) in SAA1 with change in sit-to-stand test, colored by wasting phenotype. **C-G:** Correlations of MS-measured SAA1 and SAA2 with hemoglobin (C), albumin (D), and neutrophils (E) at diagnosis (colored by CAC status), and with albumin (F) and CRP (G) after first-line treatment (colored by wasting phenotype). **H-I:** Kaplan-Meier overall survival in patients with pancreatic cancer by pre-treatment IL-6 (H) and CRP (I) tertile. Log-rank test. Numbers at risk are shown below each plot. SAA1=serum amyloid A1. SAA2=serum amyloid A2. Log2 intensity=log2-transformed concentration. nonCAC=no cachexia. preCAC=pre-cachexia. CAC=cachexia. Preserved=no wasting. FW=fat wasting. MW=muscle wasting without weight loss. MW+=muscle wasting with weight loss. CRP=C-reactive protein. HbA1c=percentage of hemoglobin that is glycated. Tau=Kendall’s tau, a non-parametric rank-correlation coefficient.

**Table S1.**
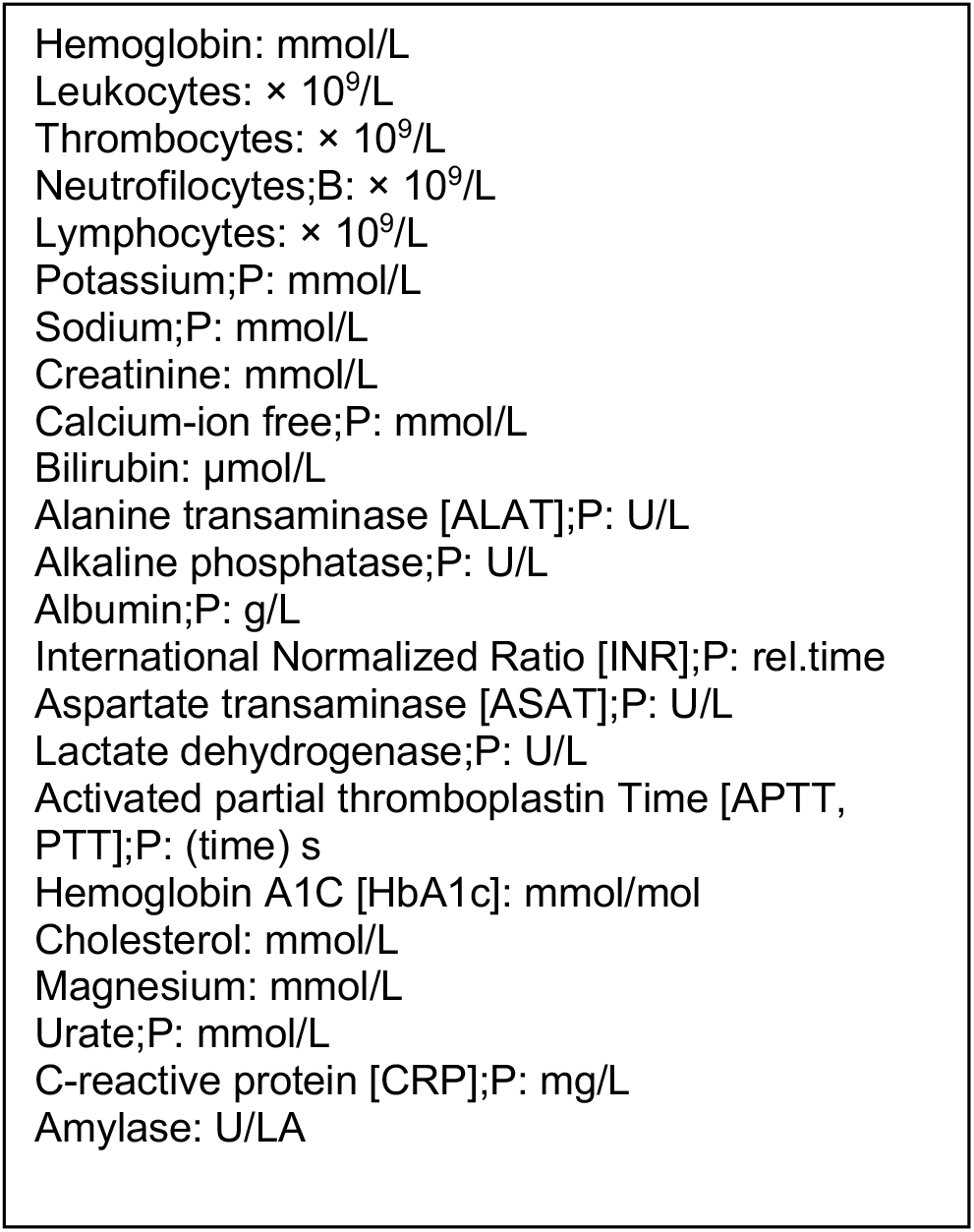
Blood samples (non-fasting) in the oncological outpatient clinic. Biochemical variables used in the selection workflow and for correlation analyses.

## METHODS

### Subjects

Patients were included within the first week after being diagnosed with advanced stage NSCLC at the Dept. of Oncology, Copenhagen University Hospital - Rigshospitalet from April 2022 to November 2023. Inclusion criteria were: (1) age > 18 years, (2) pathologically confirmed NSCLC, TNM stage IIIb/IV not eligible to concurrent chemo/radiation therapy as primary treatment, (3) referred for first-line palliative systemic anticancer therapy (4) having a staging/baseline CT scanning of the chest and abdomen within 4 weeks of initiation of treatment, or a baseline CT planned within the first week of treatment, (5) being in Eastern Cooperative Oncology Group Performance Score (PS) 0-2, (5) having signed the informed consent form to the study. Exclusion criteria were: (1) any other known malignancy requiring active treatment, (2) palliative radiotherapy as primary treatment, (3) PS >2, (4) physical disabilities excluding physical testing, (5) inability to understand Danish or (6) understand scoring systems/patient-reported outcome measures.

Patients were classified at the time of diagnosis, according to self-reported weight loss over 6 months prior to their diagnoses as: CAC (more than 5% loss of stable body weight over the past 6 months or a BMI less than 20 kg/m² and ongoing weight loss of more than 2% or sarcopenia and ongoing weight loss of more than 2% but have not entered the refractory stage)^9,25^; preCAC (1-5% weight loss); or nonCAC (increase in weight or up to 1% weight loss). Sarcopenia was defined as L3 CT-derived skeletal muscle index (SMI) (cm^2^/m^2^) < 43/41 for normal or underweight men/women and <53/41 for overweight and obese men/women)^9,25^.

Plasma samples from a clinical trial investigating IL-6 blockade in patients with locally advanced or metastatic pancreatic cancer were obtained. This randomized phase-II trial (ClinicalTrials.gov identifier: <u>NCT02767557</u>) compared the efficacy of tocilizumab, a humanized anti–IL-6 receptor antibody^18^. Patients received nab-Paclitaxel+Gemcitabine with or without tocilizumab (Table 2).

Additionally, 14 plasma samples from age and sex matched individuals from in-house biobank.

### Protocol, approvals and handling of data

The study was carried out in accordance with the Helsinki Declaration, approved by the Regional Scientific Ethical Committee of the Capital Region of Denmark (H-21035808), and registered at www.clinicaltrials.gov (NCT05307367). All clinical data was stored in a Redcap^71,75^ server operated by the Capital Region of Denmark and in agreement with the European Union’s General Data Protection Regulation (GDPR), data transfer agreement Jr.nr 22019423. Approval of processing of data was given from the local institutional board (514-0685/22-3000). Witten informed consent was obtained from each patient before participation.

### Blood sample collection

The day before the first systemic treatment was initiated (week 1) ‘at diagnosis’ blood samples were taken non-fasting in the Dept. of Oncology. In addition, blood samples were collected in week 6 and week 12 into first-line treatment. Clinical blood analyses were performed at the Department of Clinical Biochemistry, Copenhagen University Hospital – Rigshospitalet (Table S1). Plasma samples – used for MS-analyses – were collected in 9 ml EDTA tubes, kept on wet ice for <30 min, and centrifuged at 2000×g for 10 min at 4°C. Plasma was transferred to Sarstedt tubes, with 500 µl aliquots stored at −70°C.

### Body composition

In week 1, week 6 and week 12 patients’ body composition were determined via whole-body DXA-scans (DPX-IQ Lunar, enCORE v17-software, Lunar Corporation Madison, WI, USA) in Dept of Occupational therapy and Physiotherapy, Copenhagen University Hospitals - Rigshospitalet: lean body mass index (LBMI), lean mass in arms and legs known as appendicular skeletal muscle index (ASMI) and adipose tissue mass index (ATMI).

Based on the diagnostic computed tomography (CT) scan and the evaluating CT scan we determined skeletal muscle index (SMI), visceral adipose tissue index (VATI), and subcutaneous adipose tissue index (SATI). This was done by normalising the tissue area on L3 level to patient height (cm^2^/m^2^). All CT scans were done in the Depts of Radiology, Copenhagen University Hospitals - Rigshospitalet and Bispebjerg on Toshiba Prime 160 slice V0.4SP004g, Toshiba Aquillion One 320 slice V7.03GR011, GE revolution 20MW42.23 and GE Apex 20MW42.23 with 120 kV and reconstructed with 3/3 mm. Supervised by a senior consultant in radiology a research assistant handled all CT scans for further analyses. The top of the L3 vertebrae was identified. Inferior to this point, the first image clearly showing both vertebral transverse processes were selected^76,77^. From this, the selected image was pseudonymized through syngo.via (Siemens Healthineers AG). Following that, the image was exported as a DICOM file meeting the standard of being in the format 512×512 pixels. Imaging data were transferred to collaborators at Maastricht University, The Netherlands via the SURF File Sender facility allowing for application of the *Mosamatic* deep learning automated segmentation body composition software^78^. The following Hounsfield unit (HU) boundaries were used: For SAT −190 to −30 HU, for VAT −150 to −50 HU and for SM −29 to 150 HU^79^. The CT derived body compositions and their dynamics during first-line treatment allowed us to stratify the patients into 3 distinct wasting phenotypes. This is a way to capture the heterogeneity in wasting patients with cancer and CAC^16,24^. By incorporating both CT-derived loss of SMI and SATI as well as weight loss in the model and using a general cut-off on 2.5% at week 12 we wanted to reflect on the international definition of CAC and consensus of a clinical important degree of wasting per week^25^. SATI was here chosen over VATI, due to the large coefficient of variation (CV%) known from VATI data^80^. The wasting phenotypes were defined as follows: FW: loss of SATI >2.5%. MW: loss of SMI >2.5 %. MW+: loss of SMI >2.5% + weight loss >2.5%. Preserved was gain or <2.5% loss of SMI and SATI. CT scans were conducted as part of standard oncology care for disease staging at diagnosis and for evaluation of treatment response based on RECIST criteria^81^ and therefore widely available also for quantifying body composition longitudinally.

### Physical functional testing

Functional testing was conducted in week 1, week 6 and week 12 and included: Lower-limb functional strength and endurance were assessed using the 30-secound Sit-to-Stand Test^82^. Maximal Hand Grip Strength was tested by a hand-held dynamometer (Baseline BIMS Digital 5-position Grip Dynanometer, Fabrication Enterprises Inc, NY 10602, USA) best-of-three for both hands was registered and same hand was used for longitudinal comparison. Fitness Level was assessed using relative peak oxygen uptake (VO₂peak, mL·kg⁻¹·min⁻¹) which was estimated during a Cardiopulmonary Exercise Test performed on a stationary bicycle ergometer (CORTEX MetaMax 3B, MetaSoft Studio software, CORTEX Biophysik GmbH, Leipzig, Germany). The patient, wearing a heart rate monitor and a mask to measure respiration and oxygen-consumption, biked for 3 minutes at low resistance (20 watts). Resistance was then gradually increased based on a sex-specific ramp protocol until the patient could no longer continue^83^. For male patients, resistance was increased by 1 watt every 3 seconds (20 watts per minute), while for female patients, it was increased by 1 watt every 4 seconds (15 watts per minute). All patients were tested on the same bicycle and by the same 4 specially trained research physiotherapists that encouraged the patients to perform all-out before they stopped.

### Patient-reported nutritional risk

To ensure timely assessment and management of nutritional risk during 12 weeks of treatment, to minimise the impact of nutritional impact symptoms (NIS) on CAC development^23^, patients were asked to fill in the Patient-Generated Subjective Global Assessment (PG-SGA) in week 1, week 3, week 6, week 9 and week 12. An online platform linked to the Redcap server retrieved real-time patient-reported data. A PG-SGA value of ≥6 points informed the study investigator that attention from a trained oncology nurse or in-hospital dietarian team was needed^21,84^. All patients, regardless of their PG-SGA score, received standard-of-care guidance from health professionals regarding lifestyle interventions and nutrition. This was not reported.

### Sample preparation for proteomic analysis

5 μL of plasma was diluted in 45 μL lysis buffer (100 mM Tris HCl pH 8.5), and 5 μL of diluted plasma combined with 15 μL digestion buffer (100 mM Tris HCl pH 8.5), and 5 μL reduction / alkylation mix (50 mM Tcep, 200 mM CAA, 50 mM Tris). Samples were incubated for 10 min at 99°C, followed by addition of 20 μL enzyme mix (0.05 μg/μL Trypsin, 0.05 μg/μL Lys-C, 50 mM Tris pH 8.5) and incubation at 37°C for 3 h. The digest was inactivated by addition of 75 μL stop buffer (0.2% TFA). Samples were centrifuged at 4,000 g for 5 min and 25 μL of the supernatant further diluted with 75 μL stop buffer. 2.4 μL of the diluted digest were loaded onto EvoTips following the manufacturer’s instructions.

### Data acquisition by liquid chromatography–mass spectrometry LC-MS

Peptides were separated on a Pepsep 8 cm, 150 µM ID column packed with C18 beads (1.5 µm) using an Evosep ONE HPLC system. The default 60SPD method was applied, and the column temperature was maintained at 35 °C.

Upon elution, peptides were injected via an EASY-Spray source and 30-μm stainless steel emitter (Evosep) into an Orbitral Astral MS (Thermo Scientific) with FAIMS device (Thermo Scientific). FAIMS compensation voltage was set at −45V. Data was acquired in DIA mode using the MS1 in the Orbitrap at a mass resolution of 240,000 and a scan range of 380-980 m/z. The AGC target was set to 500%, and the maximum injection time was set to 10 ms. DIA scans were acquired using the Astral with a loop control set to 0.6 seconds per cycle, a maximum injection time of 5 ms, and 200 windows at 3 Th scanning from 150-2000 m/z. The HCD fragmentation normalized collision energy (NCE) was set to 25%.

### Protein identification by computational data analysis

MS files were processed using DIA-NN v.1.8.2 (beta 27) in directDIA mode. A library was predicted from the human Uniprot fasta file. Highly heuristic protein grouping and Match between runs (MBR) were enabled. Carbamidomethylation of cysteine was set as a fixed modification, while oxidation of methionine, acetylation at the protein N-terminus, and N-terminal methionine excision were set as variable modifications. The maximum missed cleavage was set to 1, and a maximum of 1 variable modification was allowed. The minimum peptide amino acid length was set to 7. Protein groups and precursors were filtered at a 1% FDR. All other settings were set as default.

### Bioinformatic processing of mass spectrometry data

All analyses were performed using RStudio (v4.3.1). A threshold of n ≥ 3 per group was applied for group-wise comparisons. Data were log2-transformed unless otherwise specified, and missing values were not imputed; therefore, analyses requiring complete datasets (e.g., principal component analysis) were conducted using only proteins without missing values. For unpaired comparisons, two-sided Welch’s t-tests were used to compute p-values. To assess the empirical p-value distribution, up to 15,000 permutations were conducted per test. The permuted p-value was calculated by dividing the number of permutation-based tests with a p-value lower than the observed p-value by the total number of permutations. If the permuted p-value exceeded the original p-value it replaced the original value. With 15,000 permutations, a p-value resolution of 0.00007 was achieved, which exceeds the significance level of all unadjusted tests. The average numerical p-value increase was 24.4 % across all comparisons, while the average reduction in significant hits was 6.4 %. All presented p-values related to the plasma proteome are shown as permutation recalibrated p-values. This was a deliberate choice given the discovery/exploratory aim of the analysis. The proteomic screen was intended to be hypothesis-generating rather than confirmatory, with candidates prioritized for in vitro validation, which provides the independent experimental support on which our conclusions rest. Over-representation analysis was performed using Homo sapiens as the background organism, with all quantified proteins used as the reference set. Gene Ontology enrichment was conducted across three categories: cellular component (GOCC), biological process (GOBP), and molecular function (GOMF). Enrichment was assessed using a hypergeometric test with Benjamini-Hochberg correction for multiple comparisons. Redundant GO terms were collapsed using a similarity threshold of 0.7. Kendall’s tau rank correlation was used for all correlation analyses.

### Plasma proteomics and workflow for target selection

Proteins were shortlisted using criteria 1 to 3 (Figure 4A) to narrow the focus to centrally regulated features considering the most divergent subgroupings at the time of inclusion (at diagnosis) and after 12 weeks. Next, we performed a network clustering analysis based on associations among shortlisted targets and selected biochemical variables measures in plasma, revealing a highly integrated network including a subpopulation of shortlisted targets. Proteins exhibiting network clustering with common biochemical measures were selected for in vitro follow up. The final proteins thus constitute the core regulation observed across subgroupings while simultaneously exhibiting a link to common biochemical markers potentially increasing the applicability in a clinical setting. Proteins of interest were those being significantly regulated with corrected p-value <0.05 and having log2FC >1.0 or < −1.0 in at least two out of three criteria. The 3 main criteria were defined: 1. CAC at diagnosis (nonCAC vs CAC). 2. Muscle wasting in week 12 (Preserved vs MW and MW+). 3 Muscle wasting in week 12 (Preserved vs MW and MW+ and FW vs. MW+). Clinical variables informing this analysis were: sit to stand-test (x/30s), hand grip strength (kg), oxygen-consumption VO2peak (fitness level; ml/min/kg), BMI (kg/m2), SMI (cm^2^/m^2^), LBMI (kg/m2), ASMI (kg/m2) and all the biochemical measures from Table S1.

### In-vitro validation of target proteins

Based on the exploration of the longitudinal plasma proteome of the patients an *in vitro* experiment using a human myoblast cell line *AB1079* was planned to test potential mediating roles of specific circulating factors for the muscle wasting observed in the patients^85^. The cell line was immortalized myoblasts from a healthy donor, provided by the Myoline facility, Institut de Myologie, Paris.

Cell culture and differentiation: AB1079 cells were cultivated in growth medium containing: 1 volume of 199 medium (Sigma-Aldrich, USA), 4 volumes DMEM (Gibco, Thermo Fisher Scientific, USA), 20 % FBS (Gibco, Thermo Fisher Scientific, USA), 50 µg/ml gentamicin (Gibco, Thermo Fisher Scientific, USA), 25 µg/ml fetuin (Sigma-Aldrich, USA), 5 ng/ml hEGF (Sigma-Aldrich, USA), 5ng/ml bFGF (Sigma-Aldrich, USA), 5 µg insulin (Actrapid, Novo Nordisk, DK) and 0.2 µg/ml dexamethasone (Sigma-Aldrich, USA). Cells were passed at a confluency below 60% and were seeded after 6-8 passages. Cells were seeded at a concentration of 20,000-25,000 cells/cm^2^ in 48 well plates (Corning, USA) pre-coated with 1% Matrigel (Corning, USA) and maintained in a humidified atmosphere of 5% CO_2_ at 37 °C. All medium volumes in the 48-well plates were 150 µl/well including seeding, differentiation and treatment. Upon 80-90 % confluency (16-24 h post seeding) differentiation was induced by replacing the growth medium with differentiation medium containing: DMEM (Gibco, Thermo Fisher Scientific, USA), 10 μg/ml of insulin (Actrapid, Novo Nordisk, Denmark) and 50 µg/ml gentamicin (Gibco, Thermo Fisher Scientific, USA).

Treatment and imaging: All recombinant proteins had a purity>95%, besides CLN5 which had a purity>90%. On day 2 of differentiation, myotubes were treated by adding treatment medium, containing: Differentiation medium, 0.1 % BSA (Thermo Fisher Scientific, USA) and 1µg/ml recombinant protein. Treatment medium was changed every 24 hours with a total of 3 treatment days. For the follow-up dose response experiment, treatment was initiated on day 5. Myotubes were treated with one dose of treatment medium containing SAA1 (1, 10 or 20 µg/ml)-Cells were imaged on day 5 or 8 of differentiation after 72 hours of treatment, using a brightfield microscope (CKX53, Olympus, Tokyo, Japan) equipped with a 10x objective with a numerical aperture of 0.25. 3 images were acquired per well for analysis of myotube width. After imaging, cells were fixed using 4% paraformaldehyde (Thermo Fisher Scientific, USA) and kept at 4 degrees Celsius until further processing. Myotube width and length was assessed using the ImageJ/FIJI software (V2.14.0/1.54f, https://imagej.net)^86^. For the initial screening, three biological replicates from three independent experiments were obtained by calculating the average from all technical replicates per condition. For the follow-up dose response, only one experiment has been analyzed thus far (ongoing work). For representative images, cells were stained with Phalloidin (Invitrogen, Massachusets, USA), DAPI (Invitrogen, Massachusets) and WGA (Invitrogen, Massachusets, USA), and mounted with fluoromount G (Invitrogen, Massachusets, USA). Images were acquired using a spinning disc confocal microscope (CSU-X1, Nikon, Japan) equipped with a 20x objective with a numerical aperture of 0.8, using Zeiss Zen software (Blue 2012 edition, Zeiss, Germany).

### Generation of human micromuscles

Human skeletal micromuscles were generated as previously described (PMID: 30527761). 44,000 human skeletal muscle myoblasts (Lonza-Lot24TL135710) were mixed in a hydrogel containing 3.3 mg/ml collagen I and 22% (v/v) Matrigel (In Vitro Technologies). The bovine acid-solubilized collagen I (Devro) was first salt balanced, and pH neutralized using 10X DMEM (ThermoFisher Scientific) and 0.1 M NaOH, respectively, prior to mixing with Matrigel and then combined with the myoblasts. The mixture was prepared on ice and pipetted into the micromuscle cell-culture inserts. The mixture was allowed to gel for 30mins at 37°C, prior to addition of growth media (1:1 mixture of DMEM:MCDB (ThermoFisher Scientific) supplemented with 4% B-27 supplement, 1% penicillin-streptomycin (P/S, ThermoFisher Scientific), 10 ng/ml FGF-2, and 100 ng/ml PDGF-BB (Invitro Technologies). After 1 day, medium was replaced with differentiation medium containing MEMα (ThermoFisher Scientific) with 1% P/S (ThermoFisher Scientific), 1% B-27 supplement (ThermoFisher Scientific), 33 μg/ml aprotinin (MedChemExpress) with 10 μM DAPT (Stem Cell Technologies) and 1 μM Dabrafenib (Stem Cell Technologies) to induce myoblast differentiation. After 7 days of differentiation, micromuscles were cultured in maintenance media consisting of MEM α (ThermoFisher Scientific) with 1% P/S (ThermoFisher Scientific), 33 μg/ml aprotinin (MedChemExpress) and 1% B-27 supplement (ThermoFisher Scientific). On day 12, micromuscles were treated with 10µg/ml Human Apo-SAA1 Recombinant Protein (PeproTech, ThermoFisher Scientific) or 0.1%BSA in PBS control.

### Analysis of force-producing capacity of human skeletal micromuscles

Human micromuscles were electrically stimulated at 1 and 20 Hz with 5ms square pulses with 20 mA current using a Digital Stimulator (Panlab/Harvard Apparatus) as previously performed (PMID: 30527761). During stimulation, an inverted high content imager (Leica DMi8) was used to capture a 5s time-lapse of each micromuscle contracting in real time at 37°C. Contractility was assessed at day 12 and day 15, and analysis was performed by Tempo.ai (Dynomics Inc).

### Murine hepatocyte isolation and cytokine treatment

Mouse hepatocytes were isolated as described previously^87^ with some modifications. Briefly, the liver was perfused with washing buffer (HBSS without calcium or magnesium + 10 mM HEPES and 200 µM EGTA (Gibco, Waltham, USA, 14170112)) through the vena cava at a flow rate of 3mL/min. The vena porta was cut to allow drainage. This was followed by perfusion of digestion buffer containing collagenase from Clostridium histolyticum (Sigma, St. Louis, C5138) in HBSS with calcium and magnesium (Gibco, Waltham, USA, 24020117). The liver was then excised and transferred to a dish containing ice-cold plating medium (Medium-199 (Gibco, Waltham, USA, 31150-022) + 10% FBS, 1% (v/v) PS), and the gallbladder was removed. The resulting cell suspension was filtered through a 100 µm filter, centrifuged, and cells washed in plating medium. Viable hepatocytes were purified by centrifugation in 90% Percoll (Sigma, St. Louis, USA, P4937) in PBS. Live cells were washed in plating medium once more before being passed through a 70 µm filter. Finally, the cells were diluted in plating medium supplemented with 0.1 mM dexamethasone (Sigma, St. Louis, USA, D4902) and 1 nM insulin (Sigma, St. Louis, USA, I9278). Hepatocytes were seeded onto collagen-coated plates at a density of 500,000 cells/mL and allowed to attach for 2-4 hours before changing to serum-free M199 culture medium for an hour before treatment with cytokines. Cells were treated with IL-1b, TNF-a and IL-6 (R&D Systems, Minneapolis, USA, 401-ML-010; 401-MT-010;406-ML-005 respectively) at the doses stated. The lowest and highest dose from individual treatments was used as the low and high combination dose, respectively.

### ELISA analysis of conditioned medium

Conditioned medium was tested. Mouse Serum Amyloid A (SAA) Quantikine ELISA kit (R&D Systems, Minneapolis, USA, MSAA00). Measurements were made according to the manufacturer’s instructions.

### Immunoblotting

Protein concentrations in lysates were measured with the bicinchoninic acid (BCA) assay (Pierce), using bovine serum albumin (BSA) standards. Samples for immunoblotting were prepared in 6× sample buffer (340 mM Tris, pH 6.8; 225 mM DTT; 11% (w/v) SDS; 20% (v/v) glycerol; 0.05% (w/v) bromophenol blue). Phosphorylated and total protein levels were measured by standard immunoblotting, with equal amounts of protein loaded per lane. Proteins were separated by SDS-PAGE on 10% gels and transferred to polyvinylidene difluoride (PVDF) membranes (Immobilon Transfer Membrane; Millipore) by semi-dry transfer. Membranes were blocked for 15 min at room temperature in Tris-buffered saline containing 0.05% Tween 20 (TBST) and 2% (w/v) skim milk, then incubated overnight at 4 °C with primary antibody. Membranes were then incubated for 1 h at room temperature with a species-specific horseradish peroxidase (HRP)-conjugated secondary antibody (Jackson ImmunoResearch). Bands were visualized with enhanced chemiluminescence (ECL+; Amersham Biosciences) on a ChemiDoc™ MP Imaging System (Bio-Rad). Densitometry was performed in Image Lab™ Software, version 4.0 (Bio-Rad; RRID: SCR_014210). Coomassie Brilliant Blue staining was used as the loading control.

### Preclinical cachexia-associated PDAC and RNAseq of liver tissue

Animal and cell culture procedures followed methods described previously in a larger, whole-body study of murine PDAC cachexia^88^. Briefly, C57BL/6J mice were purchased from Jackson Lab and acclimated to the facilities for two weeks before being randomized into experimental groups. Mice were euthanized every three days, starting on D6. Sham mice were euthanized on day 18. Twelve-week-old male mice were anesthetized under isoflurane and subcutaneously injected with Ethiqua Extended-Release 3.25mg/kg immediately before surgery and 24h from the first injection. The spleen and pancreas were gently laid back into the abdominal cavity using ring-tip forceps. The pancreas was injected with 5×10^4^ KPC32908 in 40µl of sterile PBS for 20 sec. The muscle wall was closed using absorbable sutures, and the skin flap was closed with sterile wound clips. Sham mice underwent the same procedures without manipulation of the spleen or pancreas. In parallel, KPC32908 cells were cultured and treated as outlined previously described^88,89^. At tissue collection, livers were flash frozen in liquid nitrogen for bulk RNA-sequencing.

#### Liver RNA-seq data

Sequencing data used in this study were generated as part of a larger study. Briefly, RNA was isolated from ∼30mg of snap-frozen liver tissue using the miRNeasy Mini Kit (Qiagen, Hilsen, Germany, 217004). The concentration and quality of total RNA samples were assessed using an Agilent 2100 Bioanalyzer, and an RNA Integrity Number (RIN) of eight or higher was required to pass the quality control. RNA was isolated from 4-5 mice/time point. Library preparation and sequencing were completed at the Massively Parallel Sequencing Shared Resource at Oregon Health and Science University using the Illumina NovoSeq6000. 150ng of total RNA was used for each sample. RNA sequencing, cDNA library preparation, sequencing, and filtering details were performed as previously described^88^.

### ELISA analysis of plasma samples from patients with pancreatic cancer treated with Tocilizumab and patients with non-small cell lung cancer

Plasma samples from a subset of patients with non-small cell lung cancer (Table 1), locally advanced or metastatic pancreatic cancer (Table 2; NCT05307367)^18^, and age and sex matched individuals were tested. Human Serum Amyloid A (SAA) Invitrogen™ ELISA Kit (Invitrogen, Massachusetts, USA, KHA0011C,) was used. Measurements were made according to the manufacturer’s instructions.

#### Statistics

All proteomic analysis was performed using the R Programming language within the R Studio environment.

Simple logistic regression with Holm’s method for p-adjustment was used to test socio-demographic and disease variables in included patients in analyses and dropped out. (Table 1). Dunnett’s multiple comparisons test was used to compare mean values of PG-SGA scores during treatment (Figure S1 A).

A fitted mixed linear regression with unstructured covariance matrix was used to test whether CAC groups predict nutritional score across treatment (Figure S1 B).

Unpaired t test with Welch’s correction was applied to test differences in hospitalization between nonCAC and preCAC/CAC groups (Figure S1 C). Fisher’s exact test was applied to test differences in treatment delaying toxicity (Figure S1 D).

ANOVA test was used to test differences across the different wasting phenotypes (Figure 3 A-O). Unpaired t-test was used to test differences between individual phenotypes when ANOVA p<0.05.

Parameters used in the statistical tests are as follow: CAC status at the time of diagnosis (CAC, preCAC and nonCAC); wasting phenotypes (Preserved, FW, MW and MW+).

Welch’s ANOVA was used to assess differences in clinical variables across wasting phenotypes at week 12. In cases where the overall test was significant, post hoc comparisons were conducted between the Preserved-group and each of the other groups (FW, MW, and MW+). Post hoc p-values were adjusted for multiple testing using the Benjamini-Hochberg procedure. Welch’s test p-value vs. permuted p-value was used to account for multiplicity in the comparisons.

SAA, IL-6, and CRP outliers were identified via Grubb’s test (p=0.0001) and removed prior to analysis. Outliers were excluded from control (n=2) and treatment (n=1) groups for correlation analyses, and only from the control group (n=2) for Kaplan-Meier analyses. Wilcoxon rank-sum tests compared SAA concentrations between nab-Paclitaxel+Gemcitabine and Tocilizumab+nab-Paclitaxel+Gemcitabine groups, while within-group differences were assessed using Wilcoxon signed-rank tests. At week 8 nab-Paclitaxel+Gemcitabine and Tocilizumab+nab-Paclitaxel+Gemcitabine groups were compared against each other and healthy controls via additional Wilcoxon tests.

For multiple testing, the p-value was adjusted using the Benjamini-Hochberg procedure. Kendall’s rank test was used to assess correlations between raw IL-6 and CRP VS. SAA values. Kaplan-Meier curves were fitted to the SAA values of the nab-Paclitaxel+Gemcitabine group pre-treatment. Here, the dataset was split into percentiles (33^rd^, 34^th^ to 65^th^ and 66^th^ percentile) corresponding to low, intermediate and high responders, respectively. Log-rank test was used to test for differences in overall survival. To determine differences in myotube width, a 2-way ANOVA was carried out using Prism. Statistics were calculated on biological replicates, shown as large circles in the corresponding figure, while individual technical replicates are displayed as smaller dots in corresponding colors.

## Resource availability

### Lead contact

Further information and requests for resources and reagents should be directed to the lead contact, Lykke Sylow.

### Materials availability

Materials are available upon request.

### Data and code availability

The mass spectrometry proteomics data will be deposited to the Perseus bioinformatics Platform for Integrative Analysis of Proteomics Data ^90^. This manuscript did not generate new code.

## Declaration of generative AI and AI-assisted technologies in the manuscript preparation process

During the preparation of this work the authors used Claude (Anthropic, Claude.com) in order to refine the language of the manuscript and to troubleshoot and identify syntax errors in the R script. After using this tool/service, the author(s) reviewed and edited the content as needed and take(s) full responsibility for the content of the published article.

